# Spatial cell fate manipulation of human pluripotent stem cells by controlling the microenvironment using photocurable hydrogel

**DOI:** 10.1101/2022.10.02.510557

**Authors:** Zhe Wang, Akira Numada, Fumi Wagai, Yusuke Oda, Masatoshi Ohgushi, Koichiro Maki, Taiji Adachi, Mototsugu Eiraku

## Abstract

Human pluripotent stem cells (hPSCs) dynamically respond to their chemical and physical microenvironment, dictating their behavior. However, conventional in vitro studies predominantly employ plastic culture wares, which offer a simplified representation of the in vivo microenvironment. Emerging evidence underscores the pivotal role of mechanical and topological cues in hPSC differentiation and maintenance. In this study, we cultured hPSCs on hydrogel substrates with spatially controlled stiffness.

The use of culture substrates that enable precise manipulation of spatial mechanical properties holds promise for better mimicking in vivo conditions and advancing tissue engineering techniques. We designed a photocurable polyethylene glycol–polyvinyl alcohol (PVA-PEG) hydrogel, allowing for spatial control of surface stiffness and geometry at a micrometer scale. This versatile hydrogel can be functionalized with various extracellular matrix (ECM) proteins. Laminin 511-functionalized PVA-PEG gel effectively supports the growth and differentiation of hPSCs.

Moreover, by spatially modulating the stiffness of the patterned gel, we achieved spatially selective cell differentiation, resulting in the generation of intricate, patterned structures.

**Summary statement:** A new hydrogel substrate enables spatial control of surface stiffness at the micrometer level, enabling local differentiation of hPSC and facilitating complex pattern formation.

## Introduction

hPSCs are intensively used in regenerative medicine and developmental biology research. Many types of cells and tissues have been generated from hPSCs, providing a powerful tool for *in vitro* study. However, the current culture system is distinct from the *in vivo* microenvironment, limiting the generation of complex structures *in vitro*. Pluripotent stem cells (PSCs) can be cultured for differentiation in two- or three-dimensional (2D or 3D) conditions. 2D culture provides a simple, reproducible, and robust culture condition, and many cell differentiation protocols have been developed in 2D. In contrast, in 3D culture, cells form aggregates that float in the culture medium or are embedded in an ECM (Gattazzo et al., 2014; Simunovic and Brivanlou, 2017). In both culture systems, stem cells are differentiated by adjusting the composition of the culture medium. 2D culture systems are subject to mechanical constraints on tissue shape because cells are cultured on a rigid culture dish. In contrast, in 3D culture systems, induced tissues are less mechanically constrained and can more freely change their shape. During development, stem cells organize into organs under relatively mild mechanical constraints through a combination of chemical and mechanical signals mediated by interactions between different tissues. (Simunovic and Brivanlou, 2017). Therefore, to induce functional tissues with appropriate shapes *in vitro*, an ideal environment allows tissue deformation with weak mechanical constraints.

Increasing evidence suggests the importance of mechanical cues on cell behavior and fate (Candiello et al., 2013; Chen et al., 2020; Keung et al., 2012; Maldonado et al., 2017; Muduli et al., 2017; Musah et al., 2014; Przybyla et al., 2016; Rosowski et al., 2015). Soft substrates and high mechanical tension enhance hPSC mesodermal differentiation (Muncie et al., 2020; Przybyla et al., 2016). Simultaneously, substrate geometry and structures are also critical to complex tissue structure generation (Karzbrun et al., 2021; Warmflash et al., 2014). During development, neighboring cells communicate, sensing the surrounding chemical and mechanical cues, and differentiate accordingly. Eventually, stem cells gain differentiated cell fate and cooperate to form organs with complex structures. Therefore, it is important to spatially control the microenvironment to generate complex cellular structures. To this end, we aimed to develop a culture substrate with adjustable mechanical properties.

Hydrogels have been used intensively in biomedical research as cell culture substrates due to their mechanical and chemical similarity with the ECM (Tibbitt and Anseth, 2009). Furthermore, hydrogels can be functionalized with different properties, such as photocurability. Photocurable hydrogels based on hyaluronic acid and gelatin have been developed (Chen et al., 2019; Tibbitt and Anseth, 2009). By controlling the spatial pattern of illumination, gels with different shapes and structures can be generated. However, these gels were used for hPSCs encapsulation, but not for attachment due to poor adhesivity on the synthetic surface of hPSCs (Tibbitt and Anseth, 2009; Virdi and Pethe, 2021). In contrast, some hydrogels allow suitable hPSC attachment, such as polyacrylamide or polydimethylsiloxane (PDMS), that have been used intensively in studies regarding substrate stiffness (Chen et al., 2019; Chen et al., 2020; Maldonado et al., 2017; Millar-Haskell et al., 2019; Perez-Puyana et al., 2020; Tibbitt and Anseth, 2009). However, they lack flexibility regarding gel structure and mechanical property control. Driven by this discussion, we developed a photocurable hydrogel with ideal hPSCs attachment.

hPSCs attach to the substrate via integrin-ECM protein interactions. Therefore, ECM protein engraftment on the hydrogel is necessary. Previously, ECM protein functionalized PVA could be used for hPSCs culture (Muduli et al., 2017). Polyvinyl alcohol (PVA), a polymer with a simple structure, can form a stable and biocompatible hydrogel upon cross-linking. We used PVA as the backbone and further functionalized PVA to have photocurability and ECM proteins to facilitate hPSCs attachment.

Here, we demonstrated a composite photocurable PVA and PEG hydrogel with adjustable stiffness and shape. Our hydrogel supports hPSCs growth and differentiation, and the differentiation tendency of hPSCs was compared with cells cultured on plastic dishes. Most importantly, we demonstrated that stem cell fate could be spatially manipulated at a micrometer level by controlling the local mechanical property of substrates.

## Results

### 1. Synthesis and characterization of photocurable PVA-methacrylate hydrogel

To develop a photocurable hydrogel with protein binding ability, we used a denatured Polyvinyl alcohol (PVA-COOH), which contains randomly integrated carboxyl groups as starting material. The carboxyl group in the PVA backbone can be activated by 1-ethyl-3-(3-dimethylaminopropyl) carbodiimide/N-hydroxysuccinimide (NHS/EDC) and further forms an amide bond with the N-terminus of the ECM protein to enable cell attachment (Fig. 1A). To obtain photocurability, a methacrylate functional group was introduced into the PVA backbone to obtain photocurable PVA-MA. Lithium phenyl-2,4,6-trimethyl-benzoyl phosphonate (LAP) (0.5%) was used as the photoinitiator (Fig. 1B).

**Fig. 1:**
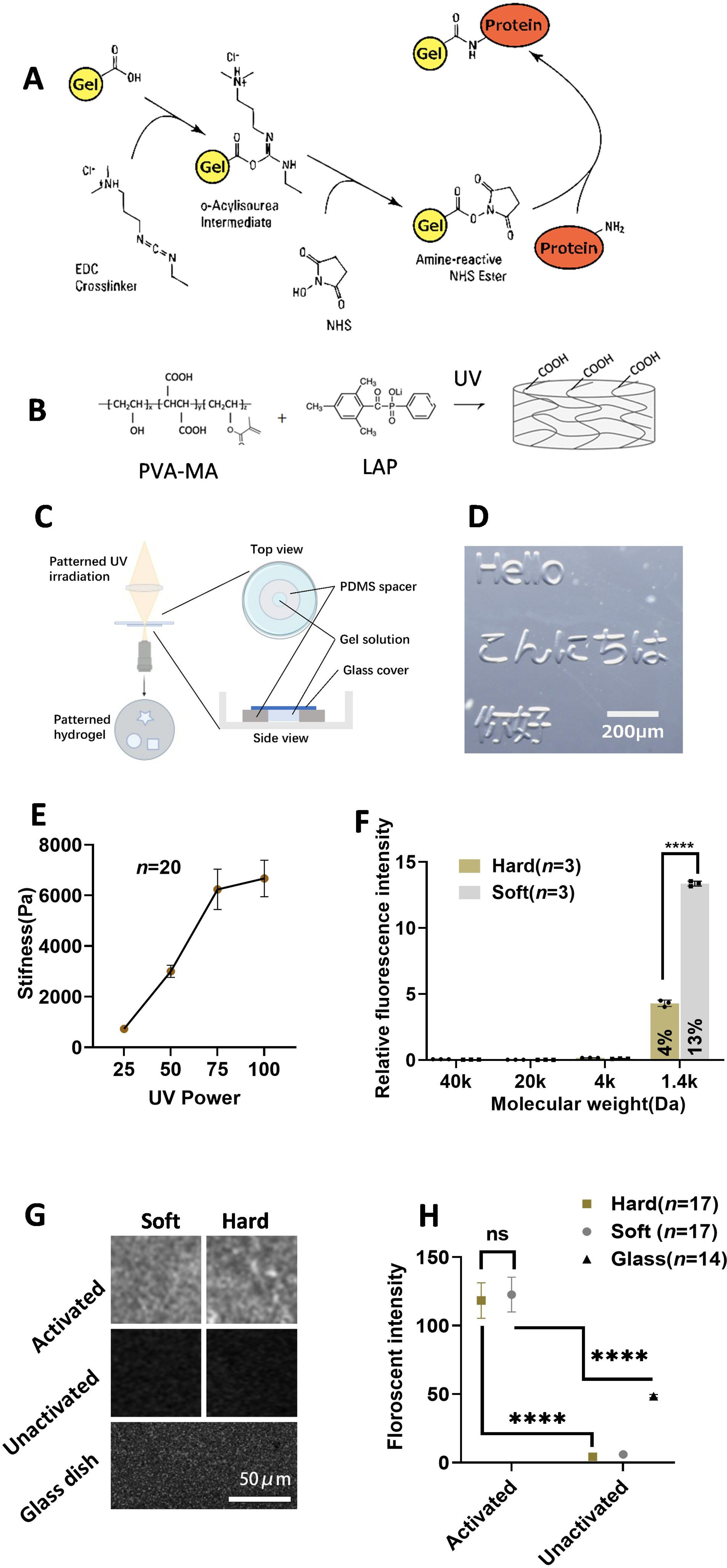
Photocurable PVA-PEG Hydrogel (A) Schematic Illustration: The hydrogel can be activated to bond with proteins using NHS/EDC.(B) Illustration: PVA-MA forms COOH-containing hydrogel upon ultraviolet (UV) irradiation.(C) Schematic Illustration: Generating patterned hydrogel.(D) Patterned Gel Generation: Patterned gel was generated using patterned UV irradiation as shown in Figure 1C.(E) Stiffness Measurement: Stiffness measured for hydrogel generated using different UV dosages. Dots represent mean values; (*n*=20). (F) Fluorescent Diffusion Experiment: Hydrogels were incubated with fluorescent molecules. Bars show the mean value of fluorescent intensity inside the gel relative to incubation solution. Black dots represent each sample. (*n*=3) (G) Pictures of Fluorescent Protein Binding Experiments: Activated and non-activated soft/hard gels, together with a glass-bottom dish, were incubated with fluorescent protein to evaluate the protein binding ability. (H) Quantification of Fluorescent Protein Binding Experiments: Fluorescent intensity was measured using fluorescence microscopy. (*n*=17 for hard, *n*=17 for soft *n*=14 for glass). Data were obtained from at least 3 biological replicates. Legends represent the mean value of each group. Scale bars: 200 µm in D, 100 µm in G. Black lines represent median values. *ns* Not significant. **** *P* < 0.0001(unpaired, two-tailed t-test). Error bars represent standard deviation.

Increasing evidence suggests the importance of the diameter of colony and substrate geometry in generating complex tissue *in vitro* (Karzbrun et al., 2021; Warmflash et al., 2014). However, current micropatterned cultures are typically constructed on glass or plastic culture ware, and the mechanical properties of cellular substrates in vitro are very different from those under in vivo conditions. Taking advantage of the flexibility of photoinitiated crosslinkability of our hydrogel, we attempted to develop a patterned culture method using our hydrogel. As seen in Fig. 2C, we generated a spacer with a round hole in the center using PDMS. The spacer was then put on the glass-bottom dish. Hydrogel prefabrication solution was added into the hole and then covered by a coverslip. We then used a direct-exposure photolithography system to spatially control the illumination pattern to initiate gelation. With this method, we were able to create various-shaped gels at the micrometer level (Fig. 1C-D). However, we encountered an issue where the PVA-MA gel alone was too soft. Previous research demonstrated that by incorporating 4-arm PEG with different molecular weights and ratios, hydrogels with stiffness ranging from approximately 500 Pa to 500 kPa could be obtained (Lee et al., 2014). Therefore, we introduced 4-arm PEG to create a composite hydrogel. We could also control the PVA-PEG composite hydrogel’s stiffness by adjusting the UV irradiation dosage. In this study, we utilized a mixture of 3% PVA and 15% 5K 4-arm PEG, as it provided us with a soft, in vivo-like stiffness and clear boundaries when generating gel patterns (Fig. 1, D-E). Consequently, we have developed a photocurable hydrogel substrate with adjustable stiffness, ranging from soft (∼700 Pa) to hard (∼6 kPa), achieved through the combination of 3% PVA and 15% 5k 4-arm PEG.

**Fig. 2:**
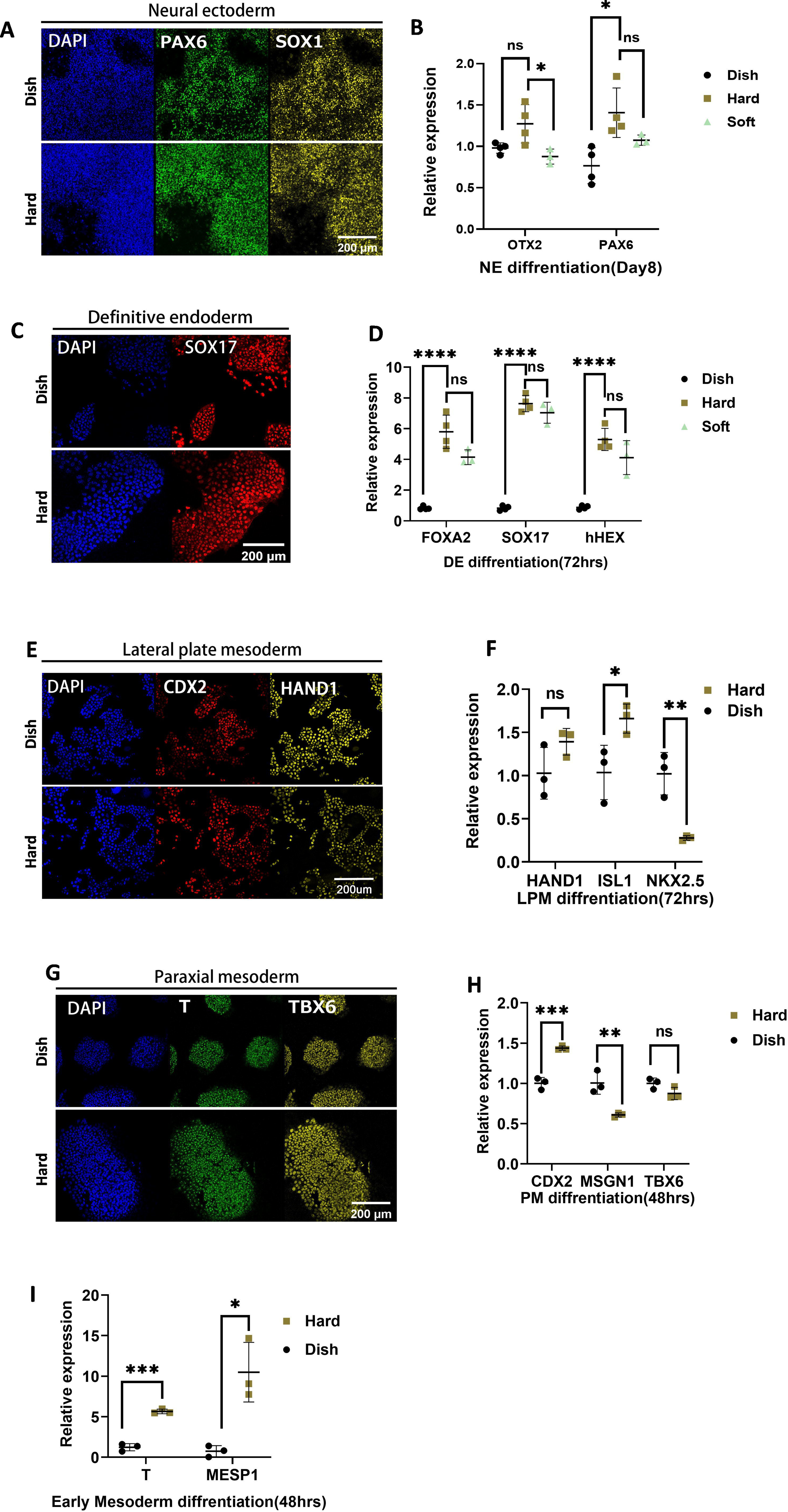
Human Pluripotent Stem Cells (hPSCs) Proliferation and YAP Localization(A) Interdivision Time: Interdivision time of hPSCs cultured on hard gel, soft gel, and conventional culture dishes. Dots represent individual cells. (*n*=75) (B) Pluripotent Marker Staining: Immunofluorescent (IF) staining of pluripotent markers in day 6 cells cultured on soft and hard gels. (C) YAP Staining: IF staining of YAP in hPSCs cultured on gels and conventional dishes. White boxes indicate 40×40 µm area magnifications of YAP staining. Data were obtained from at least 3 biological replicates. Scale bar: 100 µm. *ns* Not significant. * *P* < 0.1(unpaired, two-tailed t-test). Error bars represent standard deviation.

The crosslink density not only influences the stiffness of the hydrogel but also the mesh size of the hydrogel network. This can potentially impact cells by affecting the diffusion of proteins in the culture medium. Therefore, we investigated the diffusivity of fluorescent molecules with varying molecular weights in gels with different stiffness levels.

As illustrated in Fig. 1F and Fig. S1, molecules with a molecular weight of 4 kDa or larger were unable to diffuse within both hard and soft hydrogels. However, when using a molecule with a molecular weight of 1.4 kDa, it was capable of diffusing into both hard and soft gels, with the soft gel exhibiting higher diffusivity.

To assess the protein-binding capability of PVA-MA, we incubated both activated and non-activated PVA-MA with a fluorescent protein (Fig. 1G). Following incubation, we measured the fluorescence intensity to quantify the hydrogel’s protein-binding capacity. The activated gel exhibited a significantly higher fluorescent signal (Fig. 1H), approximately 20 times that of the non-activated hydrogel and about 2.2 times that achieved with conventional coating methods on a glass surface. This result indicates successful protein binding to our hydrogel.

### 2. PVA-PEG gels support hPSC survival and outgrowth while maintaining pluripotency

Next, we tested whether our gel supports hPSC growth. We generated two types of gels: one with a stiffness of approximately 6 kPa (hard) and another with a stiffness of around 700 Pa (soft). These gels were then activated and coated with laminin 511 as ECM proteins for hPSC culture.

To analyze cell proliferation, we developed a nucleus reporter H2B-mCherry hPSCs line that enabled us to track cell movements and conduct cell cycle analysis. H2B-mCherry-labeled cells were seeded on culture dishes or hydrogels, and live imaging was performed to monitor cell nuclear division. We found that cells cultured on the gel exhibited a similar proliferation rate compared to those cultured on glass-bottom dishes (Fig. 2A). Cells on the softer gel displayed slightly longer cell cycles, consistent with previous reports (Guo et al., 2020; Shao et al., 2015).

Immunofluorescence (IF) analysis indicated that hPSCs cultured on both hard and soft hydrogels maintained their pluripotency within one passage. This was evidenced by the expression of pluripotent markers (SOX2, OCT4, NANOG) in Day 6 cultures on the gel (Fig. 2B). These results suggest that pluripotency and proliferation of hPSCs can be sustained on hydrogels, at least within the observed time window (6 days).

It was previously known that hPSCs tend to lose nuclear-localized YAP when cultured on relatively soft substrates, a key mechano-reactive gene involved in various biological processes. Therefore, we investigated the impact of our gel on YAP localization. Cells on hard gel and conventional glass dishes exhibited clear nuclear-localized YAP, while cells on the soft gel did not (Fig. 2C, Fig. S3A-B). The hard gel exhibited lower nuclear-localized YAP than the dish (Fig.S3B). This nuclear-localized YAP can be downregulated by the Rho-kinase inhibitor Y-27632 (Fig. S3A-B). Interestingly, cells on the hard gel also displayed nuclear YAP, even though it was shown that hPSCs lost nuclear YAP on a substrate with similar stiffness (Estaras et al., 2017; Pagliari et al., 2021; Qin et al., 2016; Stronati et al., 2022). The difference from our observations is likely due to our gel’s high protein binding capacity(∼2.2 times of glass surface), which results in a dense ECM ligand (Fig. 1G-H), thus promoting nuclear YAP, as observed previously (Kechagia et al., 2023; Lee et al., 2019; Stanton et al., 2019).

### 3. Comparative analysis of hPSCs cultured on hydrogel and glass substrates by RNA sequencing

We assessed the impact of substrate stiffness on hPSCs gene expression. On Day 4, hPSCs cultures on hard gel (∼6 kPa), soft gel (∼700 Pa), and glass-bottom dishes were collected for RNA-seq analysis. Notably, pluripotent genes like SOX2, OCT3/4, and NANOG showed similar expressions in all samples (Fig. 3C), consistent with immunostaining results (Fig. 2C). We next identified differentially expressed genes (DEGs) among each sample. As a result, we found that NODAL-related genes such as NODAL, ANXA1, CER1, FOXA2, LEFTY1, LEFTY2, SMAD7, SERPINE1, and TIMP4 were upregulated in both soft and hard gels compared to culture dishes (Fig. 3D). The hard gel exhibited slightly higher endoderm gene expression compared to the soft gel (Fig. 3D). Compared to culture dishes, Gene Ontology (GO) analysis indicated that soft hydrogels enriched genes related to: Formation of the beta-catenin:TCF transactivating complex, cell morphogenesis, endoderm development, Degradation of the extracellular matrix (Fig. S2A). Hard gel enriched gene groups are: “cell morphogenesis involved in differentiation”, “endoderm development”, and “tissue morphogenesis” (Fig. S2B). Additionally, the hard gel displayed a significantly higher expression of YAP downstream genes than the soft gel, consistent with our immunofluorescence (IF) results (Fig. 2C, Fig. S2C).

**Fig. 3:**
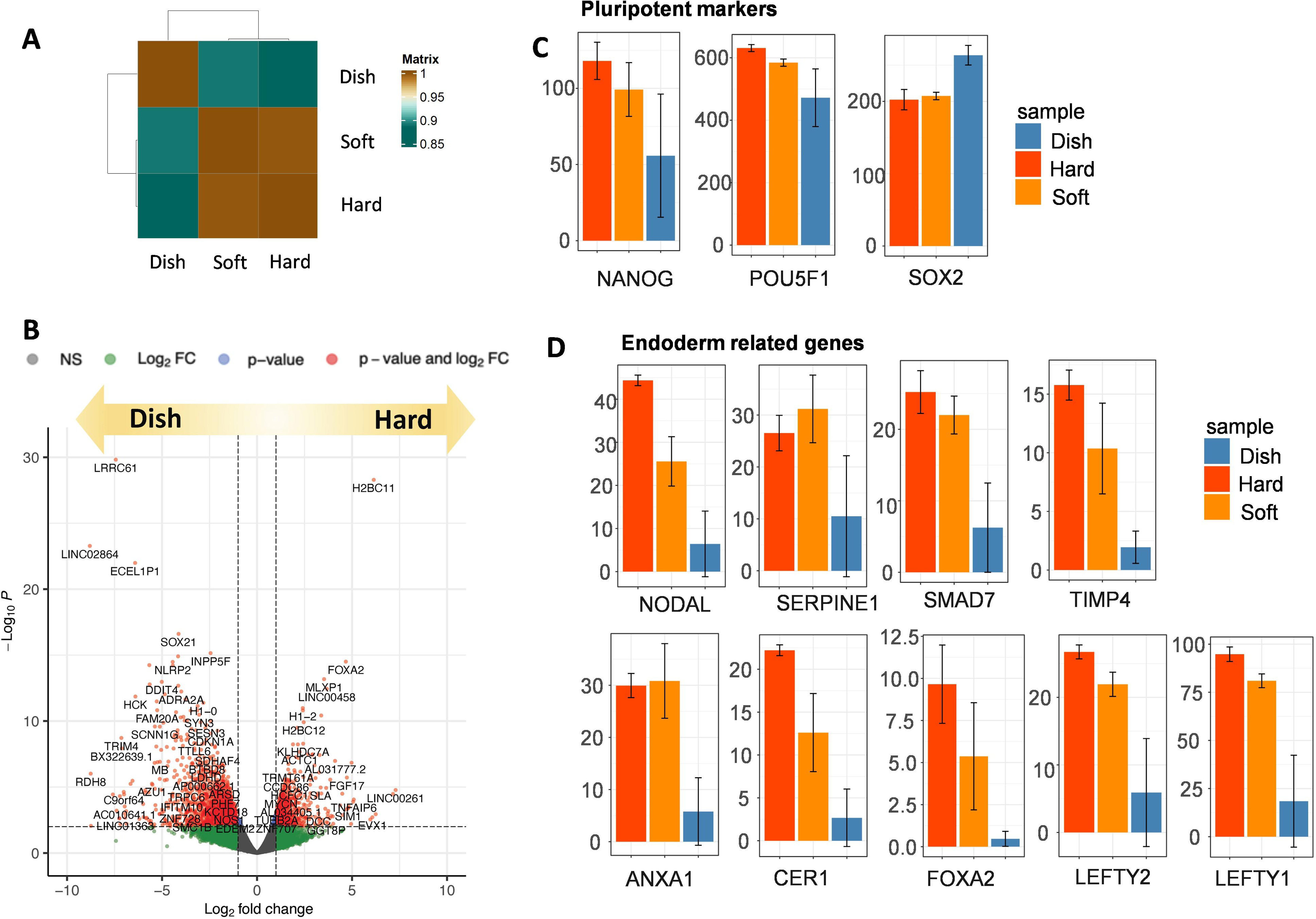
Transcriptome Analysis of hPSCs Cultured on Soft/Hard Gel and Dish. (A) Hierarchical Clustering Analysis: Hierarchical clustering analysis of mRNA expression in hPSCs cultured on soft and hard substrates compared to conventional dishes. (*n*=2), and the figure displays the mean value of the two samples. (B) Volcano Plot (Hard vs. Dish): A volcano plot representing the entire gene expression dataset for hPSCs cultured on soft gel vs. dish for 4 days. Red color indicates differentially expressed genes, and dots represent individual genes. Insignificant genes are shown in gray. (C) Pluripotent Gene Expression Based on RNA-seq Expression levels of selected genes based on RNA-seq of hPSCs cultured on soft/hard gel and dish. Expression level is shown as TPM (Transcripts Per Kilobase Million). (D) Endoderm related Gene Expression Based on RNA-seq Expression levels of selected genes based on RNA-seq of hPSCs cultured on soft/hard gel and dish. Expression level is shown as TPM (Transcripts Per Kilobase Million).

Previous reports have shown that culturing hPSCs on a 3 kPa substrate upregulates the long noncoding RNA (lncRNA) LINC00458 and LINC01356, and LINC00458 interacts with SMAD 2/3, a downstream effector of Nodal, thereby promoting endoderm differentiation (Chen et al., 2020). We noticed that these genes were also upregulated in cells cultured on the soft and hard gels (Fig. S2D).

In summary, hPSCs cultured on both soft and hard gels exhibited upregulated NODAL-related genes, possibly mediated by LINC00458 and SMAD 2/3 upregulation. Cells on the hard gel expressed higher YAP downstream genes and slightly higher but not significantly different endoderm-related genes, likely due to the preferred stiffness range for endoderm differentiation (Chen et al., 2020; Maldonado et al., 2017; Srivastava et al., 2023). Notably, both the hard and soft gels, maintained pluripotency within the observed time window, as evidenced by pluripotent gene expression (Fig. 3C).

### 4. hPSCs can be differentiated into all germ layers on PVA-PEG gel

Given the fact that our gel with different stiffness levels has varying impacts on the transcriptome, we next evaluated how our gel would affect stem cell differentiation. Cells passaged on the PS dish were collected and seeded on the hard and soft PVA-PEG gel and subsequently differentiated towards 3 germ layers following previously reported protocols (D’Amour et al., 2005; Evseenko et al., 2010; Loh et al., 2016; Surmacz et al., 2012).

For neural ectoderm (NE), hPSCs were differentiated using the previously reported dual SMAD inhibition method (Chambers et al., 2009). After 6 days of differentiation, cells on both polystyrene (PS) dish and hydrogels largely expressed the NE markers, PAX6 and OTX2, suggesting successful NE differentiation (Fig.4A-B, Fig.S4). qPCR results indicate that cells differentiated on culture dishes showed similar NE gene expression as cells differentiated on hard and soft gels. This is consistent with a previous report comparing NE differentiation on hydrogels (700 Pa, 75 kPa, 300 kPa) and culture dishes (2.28-3.28 GPa). (Fig.4B) (Keung et al., 2012; Maldonado et al., 2017).

**Fig. 4:**
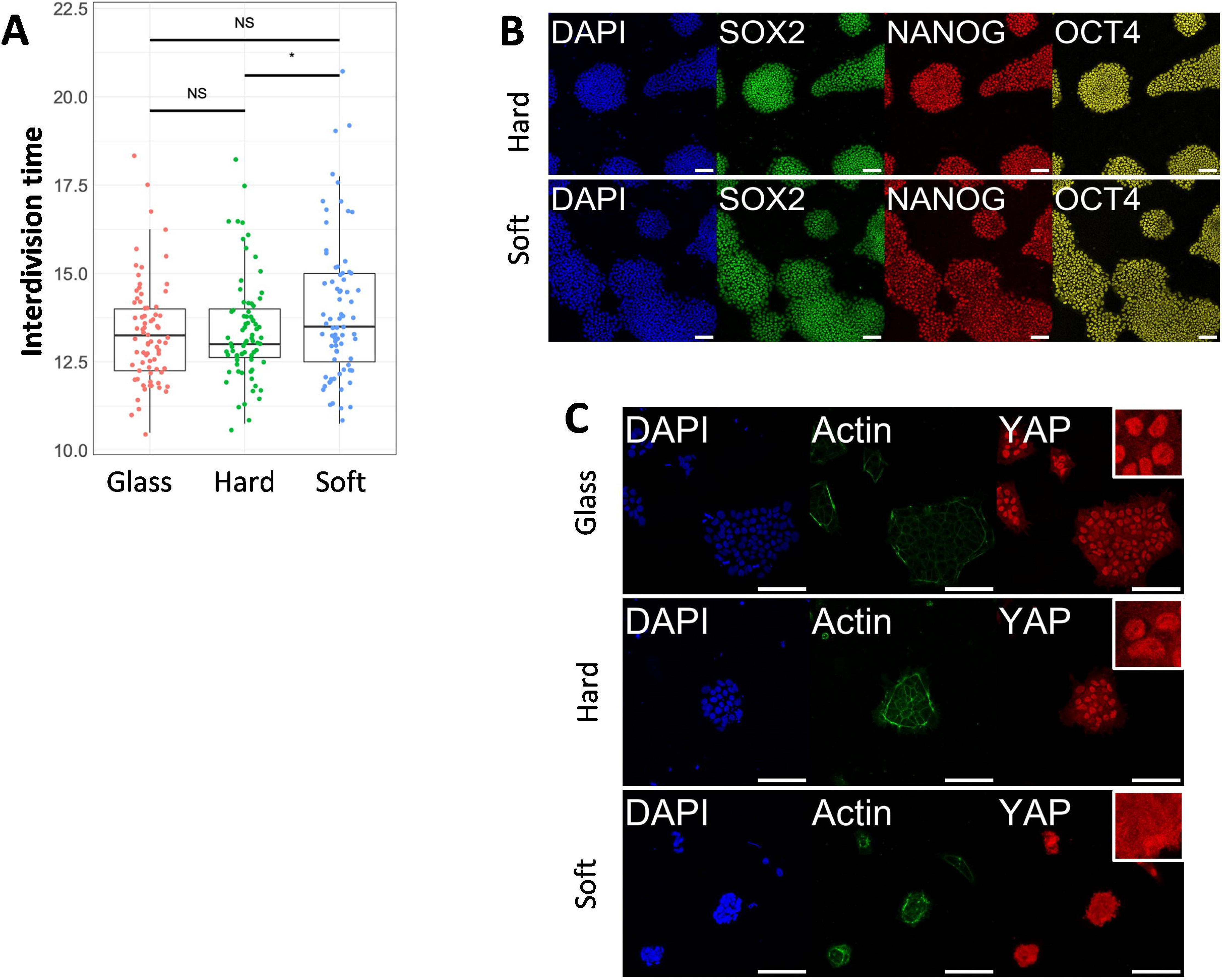
Differentiation of hPSCs on Gel (A) IF of Neural Ectoderm Differentiation (B) qPCR of Neural Ectoderm Differentiation (C) IF of Definitive Endoderm (D) qPCR of Definitive Endoderm Differentiation (E) IF of Lateral Plate Mesoderm (F) qPCR of Lateral Plate Mesoderm Differentiation (G) IF of Paraxial Mesoderm (H) qPCR of Paraxial Mesoderm Differentiation (I) qPCR of Early Mesoderm (BMP/bFGF) Differentiation. Scale bars: 200 µm. Data were obtained from at least 3 biological replicates. Each dot represents an individual biological replicate, with black lines indicating median values. Error bars represent standard deviation. Statistical significance was determined using an unpaired, two-tailed t-test. ns: Not significant **** P < 0.0001 *** P < 0.001 ** P < 0.01 * P < 0.1

For definitive endoderm (DE), cells were first differentiated to the anterior primitive streak (APS) for 24 h and then towards DE (D’Amour et al., 2005; Evseenko et al., 2010). As Fig. 4C shows, cells on both culture dishes and gels were successfully differentiated, supported by the NE marker SOX17 expression in most cells (Fig.S4). Interestingly, qPCR revealed a notably enhanced expression of APS and DE-related genes on hard and soft gels compared with the culture dish. However, there is no statistically significant difference between Soft and Hard gel (Fig. 4D).

For Mesodermal development, the paraxial mesoderm (PM) and lateral plate mesoderm (LPM) fates separate at the early stages of differentiation; therefore, we tested both.

For LPM, hESCs cells were differentiated toward mid primitive streak (MPS) for 24 h and then towards LPM for 24 h following the previously reported protocol (Loh et al., 2016). Immunostaining results suggested that the differentiation both on the gel and culture dish was successful (Fig. 4E, Fig.S4). Cells differentiated on gels expressed a higher level of *HAND1* and *ISL1* but expressed a lower level of *NKX2.5* (Fig. 4F),

For PM, hPSCs were differentiated towards APS for 24 h, and then APS was directed towards PM for 24 h (Loh et al., 2016). Although soft substrate was previously shown to enhance mesendoderm fate (Chen et al., 2020; Przybyla et al., 2016), IF and qPCR reflected similar expression of LPM markers in our experiments (Fig 4G, Fig.S4)

For both PM and LPM, qPCR revealed different differentiation efficiencies on the gel compared to conventional dishes, yet the changes remained within a similar range (less than a 2-fold change). We hypothesized that the extended differentiation processes of PM and LPM, involving multiple steps (APS to PM or MPS to LPM), could mitigate the substrate’s impact on mesoderm differentiation. Therefore, we next differentiated hPSCs into the early mesendoderm(Primitive streak) population using BMP4, as previously reported (Evseenko et al., 2010; Przybyla et al., 2016). Indeed, when compared with a conventional dish, the early mesoderm markers T and MESP1 showed a 6 and 10-fold upregulation on hard gel compared to glass dish (Fig. 4I).

These results indicate that hPSCs on our hydrogels show a similar differentiation efficiency to 3 germ layers as on PS substrates (Fig.S4), while they express higher primitive streaks gene when differentiated using BMPs on hydrogel substrates.

### 5. Manipulating cell fate of hPSCs on the patterned gel by controlling substrate stiffness

Soft substrates have been shown to enhance mesendoderm differentiation (Pagliari et al., 2021; Przybyla et al., 2016). Considering that our hydrogel enables the adjustment of stiffness in a spatially controlled manner, we were interested in exploring the potential to locally influence stem cell fate by generating soft regions within the hydrogel.

We hypothesized that if we generate locally soft patterned hydrogel, we could selectively differentiate cells on the soft part towards the early mesendoderm (primitive streak). We first generated soft and hard patterned gels and differentiated hPSCs on gels towards the mesendoderm by administrating 10 ng/mL BMP4 and bFGF (Evseenko et al., 2010; Przybyla et al., 2016) in differentiating medium (Fig. S5A). After 24 h of differentiation, cells on the soft gel were differentiated to a mesendoderm fate which was indicated by the expression of mesendodermal and primitive streak marker T. Most cells on hard gels do not express T, except a few cells on the edge (Fig. S5B), probably due to weaker cell-cell junction and higher tension (Muncie et al., 2020; Przybyla et al., 2016). These results indicate that it is possible to manipulate stem cell fate by adjusting the stiffness of the hydrogel. Since the stiffness of the gel is easily controlled by UV dosage, we next generated a hydrogel with locally different stiffness using a patterned irradiating system (Fig. 5A, Fig. S5C). We controlled the stiffness of the gel using UV dosage and created a hydrogel with locally varying stiffness through a patterned irradiation system. We confirmed the stiffness distribution in the gel pattern, which revealed a clear boundary between soft and hard areas (Fig. 5A-B).

**Fig. 5:**
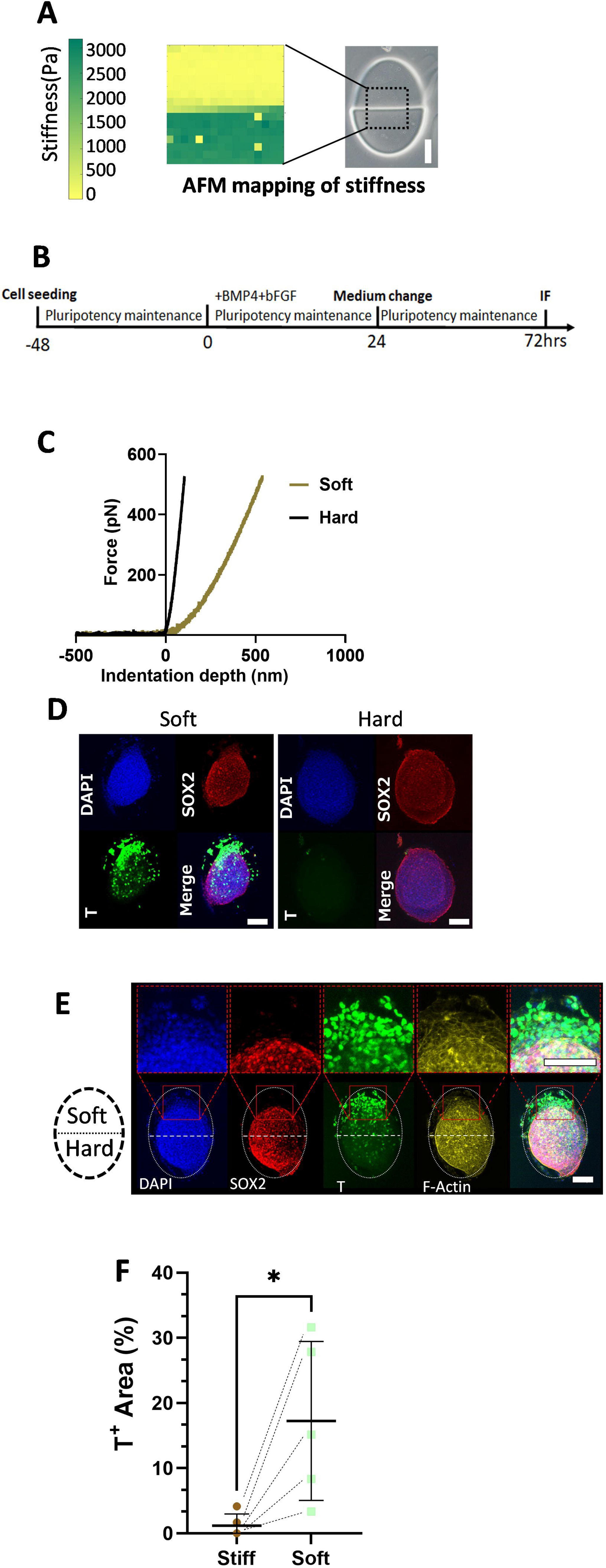
Spatial Control of Hydrogel Stiffness and Stem Cell Fate(A) Patterned Gel and AFM Stiffness Mapping: Presentation of patterned gel and a heatmap illustrating the surface stiffness of the gel measured by Atomic Force Microscope (AFM). The dotted box indicates the measured region of the hydrogel pattern. (B) Force curve of the median value of soft and hard gel acquired using AFM. The vertical axis represents the applied force (pN), while the horizontal axis represents the indentation depth(nm). The curve depicts the interaction forces between the AFM tip and the sample surface as the tip approaches the sample. (C) Schematic Illustration of Differentiation Method: A schematic illustration depicting the differentiation method. (D) IF Staining (Soft vs. Hard Gel): Immunofluorescent staining of cells differentiated on soft and hard gels using the method shown in Figure 5C. (E) IF Staining (Partially Soft Gel): Immunofluorescent staining of cells differentiated on a partially soft gel using the method shown in Figure 5C. The white dotted oval line indicates the whole gel pattern, with the upper half being soft and the lower half being hard. Red boxes show magnified views of the indicated regions. Scale bars: 100 µm. (F) Quantification of IF for Cells Differentiated on Partially Soft Gel using Pulse Differentiation: Data is represented by T+ Area/Gel surface area using Maximum projection of IF image. Dashed line-connected dots represent soft and hard parts from the same patterned gel. Data were obtained from at least 3 biological replicates. Black lines represent median values. Error bars represent standard deviation. * P < 0.05 (Paired t-test, one-tailed).

We seeded hPSCs on a patterned hydrogel, with one half being hard and the other half soft. After 24 hours of differentiation towards early mesendoderm, cells in the hard areas remained pluripotent, as indicated by positive SOX2 and negative T assays. Conversely, cells in the softer areas exhibited mesendodermal fate, as shown by their T expression and the absence of SOX2 expression (Fig. S5C). We calculated the number of T^+^ cells per area based on the IF image, as demonstrated in Fig. S5D, and found that cells on the softer gel pattern had a significantly higher (∼200 times) rate of T^+^ cells. Figure S5E illustrates the distribution of T^+^ cells per area in both soft and hard areas.

Despite the significant difference in T^+^ cell number after 24 h of differentiation, we found that cells on both the hard and soft areas became T^+^ after 48hr differentiation in the presence of BMP4 and bFGF. Therefore, we next tested if it is possible to maintain the patterned differentiation by administering a pulse of morphogen stimulation. hPSCs were seeded on hydrogel patterns and cultured in the pluripotency maintenance medium. BMP4 and bFGF (10 ng/mL each) were directly added into the maintenance medium for 24 h and then withdrawn by medium change (Fig. 5C). The cells were cultured in a maintenance medium for another 48 h. As shown in Fig. 5D, cells on the soft gel predominantly express T markers, while few or no T+ cells are observed on hard gels. Notably, on patterned gels, T+ cells are primarily localized in the soft areas, and these cells exhibit a loss of SOX2 expression, as shown in Fig. 5E-F. These data suggest that we could selectively control cell differentiation by spatially controlling the hydrogel stiffness.

We next investigated how soft gels promote early mesoderm differentiation. During gastrulation, BMP4 triggers downstream Wnt signaling, inducing primitive streak formation (Martyn et al., 2018). RNA-seq analysis also implies that enhanced β-catenin formation on the soft gel substrate (Fig. S2A), a key component of the Wnt signaling pathway. This aligns with a prior study (Przybyla et al., 2016) where cells cultured on a softer (400Pa) gel exhibited stable adherens junctions, increased β-catenin accumulation, and inhibited β-catenin degradation. Conversely, β-catenin was destabilized and degraded in cells on a harder (60kPa) gel. Consequently, cells on the softer gel exhibited heightened Wnt signaling sensitivity to BMP4-induced differentiation. To investigate the role of Wnt/β-catenin signaling, we conducted perturbation experiments. Initially, we differentiated cells on locally soft and hard gel patterns with BMP and bFGF in the presence of 2.5 µM IWP2, an inhibitor of Wnt secretion. This completely abolished differentiation, even when extended for 48 hours (Fig 6A).

**Fig. 6:**
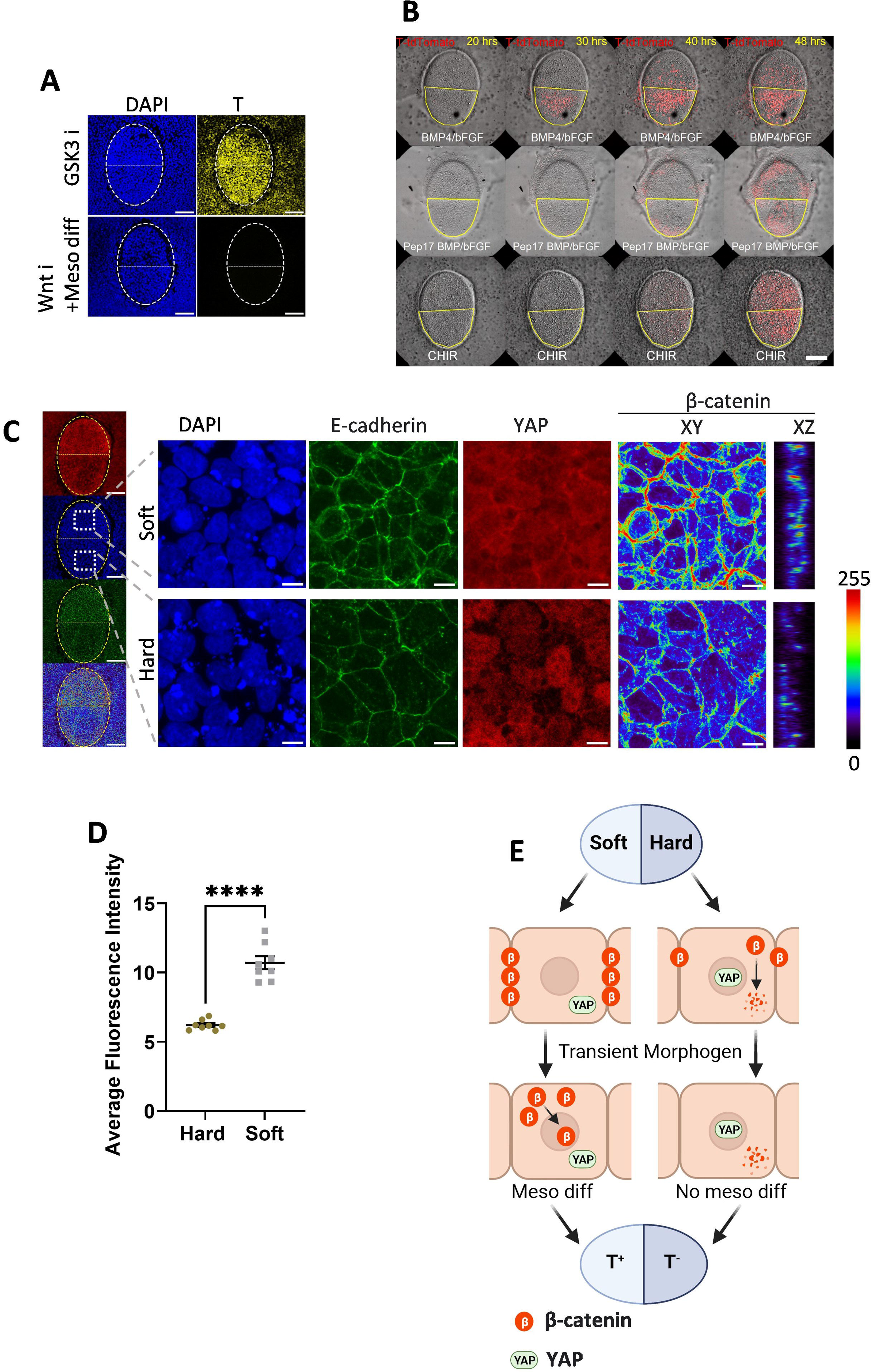
Speculated Mechanisms of Stem Cell Fate Control(A) IF Staining (GSK3 Inhibition vs. BMP/bFGF with Wnt Inhibition): Immunofluorescent staining of cells differentiated using GSK3 inhibition or by BMP/bFGF together with Wnt inhibition on a partially soft gel. The white dotted oval line indicates the whole gel pattern, with the upper half being soft and the lower half being hard. Scale bars: 100 µm. (B) Live Imaging (T-tdTomato Reporter Cell): Screenshot of live imaging of T-tdTomato reporter cells differentiated on a patterned gel using BMP/bFGF with or CHIR (Movie 1) and using BMP/bFGF in the presence of Peptide17 (Movie 2). The red color represents the tdTomato signal. The yellow line circles the soft part of the gel. Scale bars: 100 µm. (C) IF Staining (Before Differentiation): Immunofluorescent staining of hPSCs on a partially soft gel pattern before differentiation. The yellow dotted oval line indicates the whole gel pattern, with the upper half being soft and the lower half being hard. The white box indicates a magnified view of the indicated region. The β-catenin channel is shown in XY and XZ section views as a heat map, with colors representing fluorescent intensity. Scale bars: 100 µm in the whole gel view and 10 µm in the magnified view. (D) Quantification of β-catenin Fluorescent Intensity: Quantification of β-catenin fluorescent intensity based on Z sections of IF staining of the soft and hard regions on the same gel pattern. Each dot represents an individual measurement. (*n*=8). Data were obtained from at least 3 biological replicates. The black line shows the mean value. **** *P* < 0.0001 (unpaired, two-tailed t-test). Error bars represent standard deviation. (E) Proposed Mechanism of Spatial Cell Fate Control: On the soft gel, β-catenin formation is enhanced, and YAP nuclear localization is inhibited, promoting sensitivity to Wnt signaling. On the hard region, cells have nuclear-localized YAP, which negatively impacts Wnt signaling. Together with downregulated β-catenin compared to the soft part, this leads to decreased mesoderm differentiation compared to the soft region.

Conversely, differentiation occurred in both hard and soft areas when we employed the GSK3 inhibitor CHIR99201 to inhibit β-catenin degradation. This differentiation occurred without a significant time difference between soft and hard regions, unlike when differentiated with BMP (Fig. 6A-B, Fig. S6A-C Movie 1.). These results suggest that downstream Wnt/β-catenin signaling is intimately involved in the different stem cell fates induced by BMP4/FGF2 observed on hydrogels with different stiffness patterns.

As shown above, we observed a downregulation of YAP nuclear trafficking on softer gels (Fig 2C). YAP serves as a major mechano-responsive gene and has been shown to influence hESCs differentiation into various germ layers, including mesoderm. Numerous studies have reported that YAP signaling activation has a negative impact on SMAD2/3 and Wnt signaling, thereby negatively regulating mesoderm differentiation (Estaras et al., 2017; Pagliari et al., 2021; Qin et al., 2016; Srivastava et al., 2023; Stronati et al., 2022). Conversely, inhibiting YAP nuclear trafficking promotes Wnt signaling and triggers gastrulation-like differentiation (Srivastava et al., 2023). Therefore, we next investigated the YAP and β-catenin expression prior differentiation. As shown in Fig. 6E, on the hard part of the patterned gel, YAP still exhibited visible nuclear localization, while cells on the soft part displayed the opposite pattern (Fig. 6C). This aligns with the results we obtained on the non-patterned gel (Fig. 2C). Similarly, soft gel also showed stronger β-catenin staining(Fig. 2C), consistent with previous reports, a 400 Pa substrate promoted β-catenin accumulation and downregulated degradation (Przybyla et al., 2016). When differentiated with BMP4 and bFGF, this difference in β-catenin expression was still maintained (Fig. S7A-B). However, when β-catenin degradation was inhibited using the GSK3 inhibitor CHIR99201, the differences in β-catenin expression between the soft and hard regions, as well as the expression differences of T, were abolished (Fig. S7C-D). We next investigated whether the difference in stem cell fate could be abolished by inhibiting YAP signaling. T-tdTomato reporter cells on soft/hard patterned gel were pre-treated with 50nM YAP-TEAD Inhibitor 1 (Peptide 17) for 6 hours and further differentiated in the presence of Peptide 17. Live imaging revealed that this inhibition of YAP also abolished the expression differences of T in the hard and soft regions (Fig. 6B, Movie 2). Our results suggest that the spatial difference in stem cell fate, manipulated by local stiffness, is likely mediated by YAP and Wnt/β-catenin signaling. In the hard region, nuclear-localized YAP and decreased β-catenin negatively affect Wnt signaling. In contrast, in the soft region, decreased YAP nuclear trafficking and upregulated β-catenin enhance sensitivity to Wnt stimulation, resulting in spatially distinct cell fates (Fig. 6E).

## Discussion

We developed a photocurable hydrogel for hPSCs culture. We have also shown that spatially patterned illumination of UV light using a digital mirror device could control the shape and the local stiffness of the patterned hydrogels. Laminin511 E8 fragment functionalized hydrogel supports hPSCs growth and maintenance of hPSC pluripotency. hPSCs can be differentiated towards all three germ layers on our developed hydrogel. We also demonstrated when cultured on the gel, NODAL-related genes are upregulated compared to PS dishes. Most importantly, by generating locally soft gels, we could selectively differentiate hPSCs towards the mesendodermal lineage and manipulate stem cell fate spatially.

To overcome the limitation of traditional 2D/3D cultures, many soft substrates have been developed for cell cultures, such as various hydrogels, electro-spun fibers, and PDMS. However, due to the poor adhesivity of hPSCs on the synthetic surface (Tibbitt and Anseth, 2009; Virdi and Pethe, 2021), only limited materials can be used for attachment culture. PA has been used for hPSCs attachment culture; however, the pluripotency cannot be maintained. Here, Laminin 511 E8 was covalently bonded to our hydrogel and provided a strong attachment for hPSCs to enable an ideal culture of hPSCs while maintaining pluripotency.

Soft substrates such as PVA, electro-spun fibers, PA gel, and PDMS have previously been used for hPSCs attachment culture (Chen et al., 2020; Hsu et al., 2018; Keung et al., 2012; Kumbar et al., 2008; Muduli et al., 2017; Muncie et al., 2020; Musah et al., 2014; Pagliari et al., 2021; Przybyla et al., 2016; Virdi and Pethe, 2021). However, these methods usually coat the whole surface of the culture dish, and the local mechanical property cannot be adjusted. Recent research suggests the importance of the stiffness and geometry constraint in differentiating complex structures (Chen et al., 2020; Karzbrun et al., 2021; Keung et al., 2012; Martyn et al., 2018; Muncie et al., 2020; Musah et al., 2014; Przybyla et al., 2016; Rosowski et al., 2015; Srivastava et al., 2023; Virdi and Pethe, 2021). Our hydrogel provided a simple and flexible platform for generating a geometry- and local stiffness-defined culture substrate.

Currently, most of the differentiation protocols are developed by adjusting the culture medium composition. However, organs are an organized structure of different lineages. Differentiation of different lineages in a homogenous culture system by only adjusting medium composition is challenging. Here, we selectively differentiate hPSCs towards the mesendoderm by adjusting local stiffness; this provided a new tool for generating complex structures and contributing to regenerative medicine and tissue engineering.

The mechanical microenvironment plays a critical role in stem cell fate determination. We noticed that when differentiating hPSCs on different stiffness substrates, the expression changes of marker genes do not appear linear with stiffness (Maldonado et al., 2017; Musah et al., 2014). It has been previously reported that in the range of 3 kPa to 1GPa, the expression of APS and mesendoderm genes (EOMES and T) and DE genes (FOXA2 and SOX17) are inversely proportional to the substrate stiffness, but a soft substrate seems to inhibit pan primitive streak gene, CDX2, and mesodermal genes, FOXC2 and NKX2.5(Chen et al., 2020). Substrate stiffness also affects the cell fate decision developmental stage specifically. PM-related genes remained at similar expression levels in our experiment and could be explained by different observed markers, time points and stiffness range.

Furthermore, even for a similar stiffness range, studies using different materials, such as electrospun nanofibers, have different results from those using hydrogels (Chen et al., 2020; Maldonado et al., 2017; Przybyla et al., 2016), indicating the culture substrate might not only affect cells by mechanotransduction but also other factors such as the substrate dimensionality and surface chemistry. The type and distribution of ECM proteins have been shown to influence stem cell mechanosensation and fate in conjunction with substrate stiffness (Kechagia et al., 2023; Lee et al., 2019; Shibata et al., 2018; Stanton et al., 2019; Wang et al., 2015; Zhang et al., 2017). In our research, to provide stable cell attachment, we used a high density of laminin 511, known for its strong cell attachment properties. Previous reports have indicated that the high density of ECM ligands on culture substrate promotes YAP nuclear trafficking. Our gel also exhibits a higher protein binding capacity compared to commercial glass dishes (Fig. 1G-H), resulting in a denser ligand density. This might explain why stem cells exhibited nuclear-localized YAP when cultured on our hard gel (6 kPa). The interplay between ECM proteins and stiffness warrants further examination in future research.

We also found that endoderm gene expression was promoted on our gel, as indicated by our RNA-seq results (Fig. 3D), consistent with previous reports (Guo et al., 2020; Keung et al., 2012) Our developed hydrogel model promotes DE differentiation, likely through the upregulation of NODAL signaling mediated by LINC00458 and SMAD 2/3 upregulation as shown before (Guo et al., 2020; Keung et al., 2012).

The mechanism of how our hydrogel, with other substrates, affects stem cell behavior is an important aspect to explore. Most studies discussing substrate stiffness used hydrogels (Chen et al., 2020; Keung et al., 2012; Maldonado et al., 2017; Muduli et al., 2017; Muncie et al., 2020; Pagliari et al., 2021; Przybyla et al., 2016; Virdi and Pethe, 2021). Hydrogels are polymer networks containing water molecules within these networks. Changing the number of cross-links also changes the mesh size and surface chemistry, thus affecting the diffusion of the soluble factors at the basal side of the cell. In our research, we have shown that our gel can only allow small soluble factors (1.4 kDa) to penetrate but not those molecules that are bigger than 4kDa, suggesting that the stem cell fate manipulation is not caused by differences of diffusible proteins such as those of TGFβ.

We’ve shown YAP nuclear trafficking downregulation and β-catenin upregulation on softer gels. Our gel enables spatial manipulation of YAP and Wnt/β-catenin signaling for stem cell fate control in the mesendoderm differentiation. Exploring the crosstalk between YAP and Wnt/β-catenin pathways, including their hierarchy, might be a focus for future research.

In summary, we have developed a novel method for the fabrication of hPSCs culture substrate. Taking advantage of the flexibility of gel geometry and stiffness, we could precisely control the geometry and spatial stiffness. By adjusting the stiffness locally, we could selectively differentiate hPSCs toward the mesoderm, providing a novel method to generate more complex tissue structures *in vitro*.

## Methods

### Synthesis and preparation of hydrogel prefabrication solution

PVA (AP-17, Japan Vinyl Acetate & Poval Co., Ltd. Mn= 80,000–90,000) was reconstituted into a 5% PVA solution by slowly adding PVA into 50–60℃ H₂O while stirring. The temperature was then increased to 90℃ while stirring for 60 min for PVA to completely dissolve. The solution was subsequently cooled to 50 ℃ while stirring. Sulfuric acid (400 µL) was added to the PVA solution dropwise while the temperature was maintained at approximately 50℃. Methacrylic anhydride (1.5 mL) (Catalog #276685, Sigma-Aldrich, St. Louis, United States) was added to the mixture dropwise while stirring until the mixture became cloudy. More sulfuric acid was added dropwise until the solution became clear. The temperature was maintained at approximately 50℃. The reaction proceeded with stirring at 50℃ overnight. The reaction product was then transferred into a dialysis tube (Standard RC Tubing 6–8 kD Catalog #**<u>15370752</u>**, Spectrum Laboratories, Inc., Göteborg, Sweden) and dialyzed at 50℃ for 7–10 times. Obtained PVA-MA solution was then lyophilized to obtain PVA-MA powder. Prefabrication solution was prepared by reconstituting PEG, PVA-MA, and LAP in PBS to a final concentration of 10%, 5%, and 0.5% respectively. The solution was then adequate and stored in dark at -20℃. Prefabrication solution was hardened whether by using a patterned UV irradiation device in all experiments.

### Activation of hydrogel and protein binding

The carboxyl groups of the prepared hydrogels were activated by incubating the hydrogel with 10 mg/ml each of NHS/EDC in MES solution at 37℃ for 2 h. The solution was replaced with Laminin 511 E8 fragment (Imatrix511, Catalog #T304, TAKARA BIO, Kusatsu, Japan) solution (30 µL in 1 mL of PBS) incubated at 37℃ overnight. The next day, the hydrogel was washed three times with PBS and used for cell culture following standard protocol.

### Fluorescent Diffusion Test

Hydrogels were prepared as described previously. They were then incubated with solutions of fluorescent molecules in PBS for 8 hours. Cross sectional Z-stack photos of gels were taken using a fluorescence inverted microscope (SP8 LIGHTNING Confocal Microscope, Leica, Wetzlar, Germany), and the fluorescence intensity was quantified using LAS X (V 1.4.5 27713, Leica, Wetzlar, Germany). Details regarding the fluorescent molecules used can be found in Table S3.

### Cell culture

The human embryonic stem cell line, KhES-1, was originally established by Kyoto University and is routinely maintained, authenticated, and tested for contamination in our laboratory(Suemori et al., 2006). All cells used in this research are below passage number 50. KhES-1 was cultured at feeder-free conditions using StemFit^®^ AK02N (Catalog #RCAK02N, TAKARA BIO, Kusatsu, Japan) with daily medium change. Cells were routinely passaged every week. On passage day, cells were washed in PBS twice and digested using 50% CTS™ TrypLE™ Select Enzyme (Thermo fisher, Catalog #A1285901) at 37°C for 5 min. CTS™ TrypLE™ Select Enzyme was then replaced with 2 mL PBS. Cells were collected by gently pipetting and centrifuging at 600g for 5 min. ROCK Inhibitor Y-27632 (Catalog #72302, STEMCELL Technologies, Vancouver, Canada) (10 M) was added during passage to improve survival and removed the next day. Laminin 511 E8 fragment (#T304, TAKARA BIO, Kusatsu, Japan) (2.5 µg/mL) was added into the medium during passage as the cell culture matrix. The use of the KhES-1 cell line is approved and performed in accordance with the human embryonic stem cell (ESC) guidelines of the Japanese government.

### Generation of H2B-mCherry cell line

PiggyBac donor vector containing H2B fusion mCherry-IRES-puromycin resistant gene under a CAG promoter, PB-CAG-H2B-mCherry-IPuroR is a kind gift from Dr.Ohgushi. pCMV-hyPBase is a kind gift from Dr. Yusa(Yusa et al., 2011). PB-CAG-H2B-mCherry-IPuroR and pCMV-hyPBase were co-transfected into KhES-1 using Fugene 6 (Promega, E269A) transfection reagent following the manufacturer’s instruction. Three days after transfection, 1 µg/mL of puromycin was added to the culture medium to eradicate none integrated cells. Single colonies with a strong H2B-mCherry fluorescence signal were picked and expanded.

### Generation of T-H2B-tdTomato Knock in cell line

T-2A-EGFP-PGK-Puro was a gift from James Thomson (Addgene plasmid # 83344; http://n2t.net/addgene:83344; RRID: Addgene_83344). T-2A-H2B-tdTomato was generated by replacing the EGFP fragment with H2B-tdTomato using the In-Fusion^®^ HD Cloning Kit (Catalog #639648, Takara, Kusatsu, Japan), following the manufacturer’s instructions, and then linearized using PCR.

A Cas9 and sgRNA all-in-one vector, containing the sgRNA sequence ACCTTCCATGTGAAGCAGCA targeting the 3’ end of the T gene, was generated using the GeneArt^®^ CRISPR Nuclease Vector Kit (Catalog #A21174, Thermo Fisher Scientific, Waltham, United States). The linear donor was then co-transfected with the Cas9 and sgRNA all-in-one vector into KhES-1 cells using the Neon™ Transfection System 100 μL Kit (Catalog #MPK10025, Thermo Fisher Scientific, Waltham, United States) following the manufacturer’s instructions.

Three days after transfection, 1 µg/mL of puromycin was added to the culture medium to eliminate non-integrated cells. Single colonies were picked and expanded, and stock was frozen at this stage. Colonies were then differentiated using 10 μM CHIR-99021(Catalog # 72052, STEMCELL Technologies, Vancouver, Canada) for 24 hours. Colonies showing co-expression of tdTomato and T were used for subsequent experiments.

Genomic PCR was employed to detect integration. The 3’ and 5’ junctions of the HDR site in integrated colonies were further confirmed by Sanger sequencing. The puromycin resistance cassette was subsequently removed from the genome by transfecting a vector containing CRE recombinase. A single colony with puromycin resistance was picked, expanded, and used for subsequent experiments. Oligonucleotides used can be found in Table S1

### Interdivision time analysis

To compare the proliferation rate of cells cultured on gel and glass, soft or hard gel coated dish was generated following the previously described protocol. H2B-EGFP cells (1.5×10^5^) were seeded into each dish and cultured following the standard protocol. Time-lapse was performed using Olympus LCV100 confocal live imaging system (Catalog # LCV100, Olympus, Tokyo, Japan) with an interval of 15 min/frame for 72 h. Cells inside similar size colonies on gel or glass from 3 dishes were manually tracked for interdivision time analysis. The time between each nuclei division was measured according to 15 min/frame, the interdivision time of each cell was tracked at least three times and the mean value was recorded.

### Cell culture on hydrogel

Glass bottom dishes (Catalog #3961-035, AGC TECHNO, Shizuoka, Japan) was coated using Bindsilane (Catalog # 10600047, GE Healthcare, Illinois, United States) following the manufacturer’s instruction to improve hydrogel binding. Dishes were then coated with MPC polymer (Catalog # Lipidure-CM5206, NOR Corporation, Tokyo, Japan) to avoid attachment of cells on the none gel part of the dish following manufacturer instructions. The prefabrication solution was thawed and briefly vortexed. A prefabrication solution (20 µL) was added to one dish. Cover glass coated with Sigmacote^®^ (Catalog # SL2-25ML, Sigma-Aldrich, St. Louis, United States) was gently put on the gel while avoiding generating bubbles to create a flat gel surface. The gel was then hardened using Desktop Maskless Lithography System (DDB-701-DL, NEOARK CORPORATION, Hachioji, Japan). For all experiments, hard gel was hardened with 100% output power UV for 10sec, and soft gel was hardened using 25% output power for 5sec. After gelation, the cover glass was gently removed, and then hydrogel was activated and functionalized using the Laminin511 E8 fragment. Cells were seeded following standard protocol. The culture medium was half replaced the next day of passage and were all replaced 2 days after passage to gradually remove Y-27632 to avoid detachment of cells. Cells were then maintained following a standard medium change schedule.

### Stiffness Measurement of Hydrogels Using Atomic Force Microscope (AFM)

In this research, stiffness was exclusively represented by the elastic moduli. The Young’s modulus of all hydrogels was measured using an AFM (NanoWizard III, BRUKER, Billerica, United States). To briefly explain, a tipless cantilever (TL-Cont) with a spring constant of 0.02 N/m and a 10 μm diameter bead attached to the tip were employed for subsequent measurements. These measurements were conducted in the Force Spectroscopy mode within a water medium. The elastic moduli presented here are Young’s moduli, determined by applying the Hertz model to force curves. Specifically, only the initial 500 nm of indentation was considered for fitting the force curves. After noise removal using NanoWizard software (V7, BRUKER, Billerica, United States), Young’s modulus was calculated and analyzed with JPKSPM Data Processing software (V 4.2.1, JPK Instruments AG, Berlin, Germany).

### Micropatterned culture on gel

In patterning culture, cells should be seeded at high density since they are cultured on top of the hydrogel patterns. Therefore, the cell seeding area was limited using a PDMS spacer. First, PDMS sheets were prepared by mixing the main agent: hardener = 9:1 for 15 min, and then pour onto a flat surface and cured by heating at 80°C for 15–20 min. The sheet was made as thin as possible for subsequent patterning culture. A hole of 6 mm diameter was made in the center of the formed PDMS sheet using a hole punch. PDMS spacer was attached to Bindsilane coated 35 mm glass-bottom dish and 20 µL of PVA-PEG prefabrication solution was added into the hole. Then Sigmacote^®^ coated coverslip was gently put on top of the gel solution. The patterned gel was then cured by irradiating patterned UV using Desktop Maskless Lithography System (DDB-701-DL, NEOARK CORPORATION, Hachioji, Japan). Gel with different stiffness was generated by giving spatially different UV exposure (100% output for 10 seconds for the hard part and 25% output for 5 seconds for the soft part). Then the coverslip was removed, and hydrogel was activated and functionalized with Laminin511 E8 fragment. hESCs (KhES-1) (1×10^6^) were collected and resuspended in 200 µL of AK02N medium with 10 µM Y27632.The suspension (20 L) was added on top of the gel pattern and incubated at 37 °C, 5% CO_2_ for 30 min to allow cells to attach. The gel was washed with PBS once to remove exceeded cells and the PDMS spacer was removed. AK02N medium (1 mL) with 10uM Y27632 was added to the dish. Half of the medium was replaced with AK02N the next day. Then Cells were cultured following standard protocol.

### Mesoderm differentiation

Mesoderm differentiation was done following previously reported method(Evseenko et al., 2010). Shortly, hESCs cultured on gel pattern was generated following protocol mentioned above. Cell culture medium was replaced with Stemline II Hematopoietic Stem Cell Expansion Medium (Catalog # S0192-500ML, Sigma-Aldrich, St. Louis, United States) or AK02N(for pulse differentiation) supplemented with 10ng each of BMP4(Catalog # H4916, Sigma-Aldrich, Tokyo, Japan) and bFGF(Catalog # 100-18B-1MG, Thermo Fisher Scientific, Waltham, USA) for 24hrs.

### Live imaging of T-H2B-tdTomato

We generated a patterned culture on a gel with an oval shape, where half of it was soft, and the other half was hard, as previously described. In brief, T-H2B-tdTomato cells cultured on the gel pattern, which had locally different stiffness, were subjected to differentiation using either 10μM CHIR or 10ng/ml each of BMP4 and bFGF for 24 hours. Subsequently, the differentiating medium was removed, and the samples were washed twice with PBS before being returned to the maintenance culture medium.

In YAP inhibition experiments, the cells were pretreated with Peptide 17 (Catalog # S8164, Selleck Chemicals, Texas, United States) for 6 hours before they were differentiated with 10ng/ml each of BMP4 and bFGF under the presence of Peptide 17. Following differentiation, the cells were subjected to live imaging.

For imaging of T expression, we utilized the Olympus LCV100 confocal live imaging system, capturing images at intervals of 60 minutes per frame for a duration of 48 hours.

### Measuring protein binding ability of PVA-MA

PVA-MA hydrogel was prepared and activated as described previously. The hydrogel was then incubated with a fluorescent 2nd antibody (Catalog # A-11003, ThermoFisher, Massachusetts, United States) following the protein binding protocol described previously. Z stack Photos of the gel surface were taken using a fluorescence inverted microscope (SP8 LIGHTNING Confocal Microscope, Leica, Wetzlar, Germany) and the fluorescence intensity was quantified using LAS X(V 1.4.5 27713, Leica, Wetzlar, Germany).

### Immunofluorescence

The culture medium was removed from the sample. Samples were then fixed by incubating in 2 mL of 4% PFA in PBS on ice for 20 min. Samples were washed in PBS three times for 5 min each. PBS was replaced with 0.3% Triton-X100/PBS and samples were incubated for 5 min at 25℃. Samples were then blocked with Blocking One (Catalog #03953-66, NACALAI TESQUE, Kyoto, Japan) for 1 h at room temperature, then incubated with primary antibody diluted in blocking one at 4°C overnight. primary antibody was Removed then samples were washed with 0.05% Tween 20/PBS for 5 min and repeated three times. Then samples were incubated with secondary antibodies and DAPI diluted in the blocking solution, 1/500 for 1 h at room temperature. Secondary antibodies were removed, and samples were washed in 0.05% Tween 20/PBS for 5 min, thrice. The samples were then observed using a microscope.

### RNA extraction and quantitative PCR

Total RNA was extracted using RNeasy Mini Kit (Catalog #74106, QIAGEN, Hilden, Germany) following the manufacturer’s instructions. cDNA was reverse transcripted from 500ng RNA using PrimeScript™ II 1st strand cDNA Synthesis Kit (Catalog # 6210A TAKARA BIO, Kusatsu, Japan) following the manufacturer’s instructions. qPCR was performed using Power™ SYBR™ Green Master Mix (Catalog #4368577, Applied Biosystems™, Massachusetts, United States). The expression level of mRNAs was calculated and normalized based on indicated housekeeping gene (*HPRT1*, *PBGD*).

### RNA-seq

Gel-coated 35 mm glass-bottom dishes were prepared as previously described. KhES-1 was seeded on the soft gel, hard gel, or uncoated glass-bottom dish and cultured for 4 days before they were collected as previously described. Total RNA was extracted using RNeasy Mini Kit (Cat#74106, QIAGEN, HILDEN, GERMANY,) following the manufacturer’s instruction. Two samples for each group, 6 samples in total were collected. Libraries were prepared from 100 ng of total RNA using the following kits, NEBNext Poly(A) mRNA Magnetic Isolation Module (Catalog # E7490, NEW ENGLAND BIOLABS, MASSACHUSETTS, UNITED STATES), NEBNext Ultra II Directional RNA Library Prep Kit (Catalog # E7760, NEW ENGLAND BIOLABS,MASSACHUSETTS, UNITED STATES), NEBNext Multiplex Oligos for Illumina (96 Unique Dual Index Primer Pairs) (Catalog # E6440, NEW ENGLAND BIOLABS,MASSACHUSETTS, UNITED STATES) following the manufacturer’s instructions. Sequencing was performed with the paired-end mode (more than 50 bp at both ends) to have at least 15 clusters per sample using the NextSeq550 NGS system.

The raw RNA-seq files generated were quality controlled using FastQC version 0.11.8 and Trimomatic version 0.39. High-quality sequence reads were then mapped and annotated with STAR version 2.7.6a against the GRCh38 reference genome. Subsequently, count data was estimated using subread version 2.0.1 and subjected to differential gene expression analysis using the Bioconductor package DESeq2 version 1.30.1 in R version 4.0.3. Differentially expressed genes (DEGs) were defined as genes with a false discovery rate (FDR) < 0.1. The Metascape database platform was utilized to enrich and analyze Gene Ontology (GO) annotations. For the TPM matrix, RSEM version 1.3.3 was used to quantify data from STAR-mapped reads(Li and Dewey, 2011).

### YAP quantification

Nucleocytoplasmic transport of YAP was quantified using the N/C intensity ratio (Kelley and Paschal, 2019) .In brief, cells cultured on hard and soft gels or on a rigid dish were subjected to immunofluorescence (IF) staining following the previously mentioned protocol. Z-stack images were acquired, and a Z-projection was generated using the Sum Slices method in ImageJ2(Rueden et al., 2017). To mitigate the impact of confluency on YAP localization, we focused on quantifying small colonies containing fewer than 20 cells.

In summary, the process involved thresholding a DAPI image for nuclear staining to create a mask defining the nuclear area. Simultaneously, the YAP-stained channel was thresholded to delineate the cytoplasmic area containing the nuclear region. The nuclear area was then subtracted from the YAP-stained image using the DAPI-generated mask. The resulting area was utilized to measure the total fluorescence intensity of the cytoplasm. The Nuclear/Cytoplasmic (N/C) ratio was calculated as follows: N/C = total intensity of the nucleus / total intensity of the cytoplasm.

### Statistical analysis

Statistical analysis was done using GraphPad Prism (V 9.3.1, GraphPad Software, La Jolla, United States). Two tail unpaired t test was used to compare the differences between 2 groups. All data were derived from a minimum of three biological replicates.

### Manuscript preparation

Manuscript was first written by the first author, ChatGPT (V 3.5, Open AI, San Francisco, US) was used to check and improve the clarity and grammar of written manuscript.

## Acknowledgments

We extend our gratitude to the Single-Cell Genome Information Analysis Core (SignAC) at WPI-ASHBi, Kyoto University, for their assistance in RNA sequencing. Special thanks to Kosuke Yusa for providing the PiggyBac vector. Our sincere appreciation goes to all lab members for their valuable suggestions and assistance, with special acknowledgment to Rio Tsutsumi and Yusuke Seto. We would also like to express our thanks to Yasuhiko TABATA and Wenxuan Yang for their assistance with frozen drying.

## Funding

*T*his work was supported by Grant-in-Aid for Scientific Research on Innovative Areas (Ministry of Education, Culture, Sports, Science, and Technology (MEXT)), Japan (16H06480 to M.E.), Grant-in-Aid for Transformative Research Areas (MEXT), Japan (23H04933 to M.E.), and Core Research for Evolutional Science and Technology (CREST, JST) (JPMJCR12W2 to M.E.), Japan Science and Technology Agency, Support for Pioneering Research Initiated by the Next Generation (Grant Number JPMJSP2110 to Z.W.), Japan Society for the Promotion of Science ,Grant-in-Aid for JSPS Research Fellow (Grant Number 22J10657 to Z.W.).

## Author contributions

Z.W., A.N., and M.E. designed the research; Z.W. and A.N. performed the research; Z.W., A.N., F.W., and Y.O. analyzed the data. M. O. made plasmid for H2B-GFP cell line. K. M. and T. A. helped and performed AFM analysis and gave feedback on the manuscript. Z.W. and M.E wrote the paper.

## Data availability statement

The data and materials supporting this research are available from the authors on reasonable request. RNA-seq data has been deposited in the SRA database and may be accessed with the accession number PRJNA1022777

## Additional information

The authors declare no competing interests.

## Notes

### Competing Interest Statement

The authors have declared no competing interest.

### Summary of Updates

2nd revision of development

## References

Candiello, J., Singh, S. S., Task, K., Kumta, P. N. and Banerjee, I. (2013). Early differentiation patterning of mouse embryonic stem cells in response to variations in alginate substrate stiffness. J Biol Eng 7, 9.

Chambers, S. M., Fasano, C. A., Papapetrou, E. P., Tomishima, M., Sadelain, M. and Studer, L. (2009). Highly efficient neural conversion of human ES and iPS cells by dual inhibition of SMAD signaling. Nat Biotechnol 27, 275-280.

Chen, G., Tang, W., Wang, X., Zhao, X., Chen, C. and Zhu, Z. (2019). Applications of Hydrogels with Special Physical Properties in Biomedicine. Polymers (Basel) 11.

Chen, Y. F., Li, Y. J., Chou, C. H., Chiew, M. Y., Huang, H. D., Ho, J. H., Chien, S. and Lee, O. K. (2020). Control of matrix stiffness promotes endodermal lineage specification by regulating SMAD2/3 via lncRNA LINC00458. Sci Adv 6, eaay0264.

D’Amour, K. A., Agulnick, A. D., Eliazer, S., Kelly, O. G., Kroon, E. and Baetge, E. E. (2005). Efficient differentiation of human embryonic stem cells to definitive endoderm. Nat Biotechnol 23, 1534-1541.

Estaras, C., Hsu, H. T., Huang, L. and Jones, K. A. (2017). YAP repression of the WNT3 gene controls hESC differentiation along the cardiac mesoderm lineage. Genes Dev 31, 2250-2263.

Evseenko, D., Zhu, Y., Schenke-Layland, K., Kuo, J., Latour, B., Ge, S., Scholes, J., Dravid, G., Li, X., MacLellan, W. R., et al. (2010). Mapping the first stages of mesoderm commitment during differentiation of human embryonic stem cells. Proc Natl Acad Sci U S A 107, 13742-13747.

Gattazzo, F., Urciuolo, A. and Bonaldo, P. (2014). Extracellular matrix: a dynamic microenvironment for stem cell niche. Biochim Biophys Acta 1840, 2506-2519.

Guo, A., Wang, B., Lyu, C., Li, W., Wu, Y., Zhu, L., Bi, R., Huang, C., Li, J. J. and Du, Y. (2020). Consistent apparent Young’s modulus of human embryonic stem cells and derived cell types stabilized by substrate stiffness regulation promotes lineage specificity maintenance. Cell Regeneration 9.

Hsu, H.-T., Estarás, C., Huang, L. and Jones, K. A. (2018). Specifying the anterior primitive streak by modulating YAP1 levels in human pluripotent stem cells. Stem cell reports 11, 1357-1364.

Karzbrun, E., Khankhel, A. H., Megale, H. C., Glasauer, S. M. K., Wyle, Y., Britton, G., Warmflash, A., Kosik, K. S., Siggia, E. D., Shraiman, B. I., et al. (2021). Human neural tube morphogenesis in vitro by geometric constraints. Nature 599, 268-272.

Kechagia, Z., Saez, P., Gomez-Gonzalez, M., Canales, B., Viswanadha, S., Zamarbide, M., Andreu, I., Koorman, T., Beedle, A. E. M., Elosegui-Artola, A., et al. (2023). The laminin-keratin link shields the nucleus from mechanical deformation and signalling. Nat Mater.

Kelley, J. B. and Paschal, B. M. (2019). Fluorescence-based quantification of nucleocytoplasmic transport. Methods 157, 106-114.

Keung, A. J., Asuri, P., Kumar, S. and Schaffer, D. V. (2012). Soft microenvironments promote the early neurogenic differentiation but not self-renewal of human pluripotent stem cells. Integrative Biology 4, 1049-1058.

Kumbar, S. G., James, R., Nukavarapu, S. P. and Laurencin, C. T. (2008). Electrospun nanofiber scaffolds: engineering soft tissues. Biomed Mater 3, 034002.

Lee, S., Stanton, A. E., Tong, X. and Yang, F. (2019). Hydrogels with enhanced protein conjugation efficiency reveal stiffness-induced YAP localization in stem cells depends on biochemical cues. Biomaterials 202, 26-34.

Lee, S., Tong, X. and Yang, F. (2014). The effects of varying poly(ethylene glycol) hydrogel crosslinking density and the crosslinking mechanism on protein accumulation in three-dimensional hydrogels. Acta Biomater 10, 4167-4174.

Li, B. and Dewey, C. N. (2011). RSEM: accurate transcript quantification from RNA-Seq data with or without a reference genome. BMC Bioinformatics 12, 323.

Loh, K. M., Chen, A., Koh, P. W., Deng, T. Z., Sinha, R., Tsai, J. M., Barkal, A. A., Shen, K. Y., Jain, R., Morganti, R. M., et al. (2016). Mapping the Pairwise Choices Leading from Pluripotency to Human Bone, Heart, and Other Mesoderm Cell Types. Cell 166, 451-467.

Maldonado, M., Luu, R. J., Ico, G., Ospina, A., Myung, D., Shih, H. P. and Nam, J. (2017). Lineage- and developmental stage-specific mechanomodulation of induced pluripotent stem cell differentiation. Stem Cell Res Ther 8, 216.

Martyn, I., Kanno, T. Y., Ruzo, A., Siggia, E. D. and Brivanlou, A. H. (2018). Self-organization of a human organizer by combined Wnt and Nodal signalling. Nature 558, 132-135.

Millar-Haskell, C. S., Dang, A. M. and Gleghorn, J. P. (2019). Coupling synthetic biology and programmable materials to construct complex tissue ecosystems. MRS Commun 9, 421-432.

Muduli, S., Chen, L. H., Li, M. P., Heish, Z. W., Liu, C. H., Kumar, S., Alarfaj, A. A., Munusamy, M. A., Benelli, G., Murugan, K., et al. (2017). Stem cell culture on polyvinyl alcohol hydrogels having different elasticity and immobilized with ECM-derived oligopeptides. J Polym Eng 37, 647-660.

Muncie, J. M., Ayad, N. M. E., Lakins, J. N., Xue, X., Fu, J. and Weaver, V. M. (2020). Mechanical Tension Promotes Formation of Gastrulation-like Nodes and Patterns Mesoderm Specification in Human Embryonic Stem Cells. Dev Cell 55, 679-694 e611.

Musah, S., Wrighton, P. J., Zaltsman, Y., Zhong, X., Zorn, S., Parlato, M. B., Hsiao, C., Palecek, S. P., Chang, Q., Murphy, W. L., et al. (2014). Substratum-induced differentiation of human pluripotent stem cells reveals the coactivator YAP is a potent regulator of neuronal specification. Proceedings of the National Academy of Sciences 111, 13805-13810.

Pagliari, S., Vinarsky, V., Martino, F., Perestrelo, A. R., Oliver De La Cruz, J., Caluori, G., Vrbsky, J., Mozetic, P., Pompeiano, A., Zancla, A., et al. (2021). YAP–TEAD1 control of cytoskeleton dynamics and intracellular tension guides human pluripotent stem cell mesoderm specification. Cell Death & Differentiation 28, 1193-1207.

Perez-Puyana, V., Jimenez-Rosado, M., Romero, A. and Guerrero, A. (2020). Polymer-Based Scaffolds for Soft-Tissue Engineering. Polymers (Basel) 12.

Przybyla, L., Lakins, J. N. and Weaver, V. M. (2016). Tissue Mechanics Orchestrate Wnt-Dependent Human Embryonic Stem Cell Differentiation. Cell Stem Cell 19, 462-475.

Qin, H., Hejna, M., Liu, Y., Percharde, M., Wossidlo, M., Blouin, L., Durruthy-Durruthy, J., Wong, P., Qi, Z., Yu, J., et al. (2016). YAP Induces Human Naive Pluripotency. Cell Rep 14, 2301-2312.

Rosowski, K. A., Mertz, A. F., Norcross, S., Dufresne, E. R. and Horsley, V. (2015). Edges of human embryonic stem cell colonies display distinct mechanical properties and differentiation potential. Sci Rep 5, 14218.

Rueden, C. T., Schindelin, J., Hiner, M. C., DeZonia, B. E., Walter, A. E., Arena, E. T. and Eliceiri, K. W. (2017). ImageJ2: ImageJ for the next generation of scientific image data. BMC Bioinformatics 18, 529.

Shao, Y., Sang, J. and Fu, J. (2015). On human pluripotent stem cell control: The rise of 3D bioengineering and mechanobiology. Biomaterials 52, 26-43.

Shibata, S., Hayashi, R., Okubo, T., Kudo, Y., Katayama, T., Ishikawa, Y., Toga, J., Yagi, E., Honma, Y., Quantock, A. J., et al. (2018). Selective Laminin-Directed Differentiation of Human Induced Pluripotent Stem Cells into Distinct Ocular Lineages. Cell Rep 25, 1668-1679 e1665.

Simunovic, M. and Brivanlou, A. H. (2017). Embryoids, organoids and gastruloids: new approaches to understanding embryogenesis. Development 144, 976-985.

Srivastava, P., Romanazzo, S., Kopecky, C., Nemec, S., Ireland, J., Molley, T. G., Lin, K., Jayathilaka, P. B., Pandzic, E., Yeola, A., et al. (2023). Defined Microenvironments Trigger In Vitro Gastrulation in Human Pluripotent Stem Cells. Adv Sci 10.

Stanton, A. E., Tong, X., Lee, S. and Yang, F. (2019). Biochemical Ligand Density Regulates Yes-Associated Protein Translocation in Stem Cells through Cytoskeletal Tension and Integrins. ACS Appl Mater Interfaces 11, 8849-8857.

Stronati, E., Giraldez, S., Huang, L., Abraham, E., McGuire, G. R., Hsu, H. T., Jones, K. A. and Estaras, C. (2022). YAP1 regulates the self-organized fate patterning of hESC-derived gastruloids. Stem Cell Reports 17, 211-220.

Suemori, H., Yasuchika, K., Hasegawa, K., Fujioka, T., Tsuneyoshi, N. and Nakatsuji, N. (2006). Efficient establishment of human embryonic stem cell lines and long-term maintenance with stable karyotype by enzymatic bulk passage. Biochem Biophys Res Commun 345, 926-932.

Surmacz, B., Fox, H., Gutteridge, A., Fish, P., Lubitz, S. and Whiting, P. (2012). Directing differentiation of human embryonic stem cells toward anterior neural ectoderm using small molecules. Stem Cells 30, 1875-1884.

Tibbitt, M. W. and Anseth, K. S. (2009). Hydrogels as extracellular matrix mimics for 3D cell culture. Biotechnol Bioeng 103, 655-663.

Virdi, J. K. and Pethe, P. (2021). Biomaterials regulate mechanosensors YAP/TAZ in stem cell growth and differentiation. Tissue Engineering and Regenerative Medicine 18, 199-215.

Wang, H., Luo, X. and Leighton, J. (2015). Extracellular Matrix and Integrins in Embryonic Stem Cell Differentiation. Biochem Insights 8, 15-21.

Warmflash, A., Sorre, B., Etoc, F., Siggia, E. D. and Brivanlou, A. H. (2014). A method to recapitulate early embryonic spatial patterning in human embryonic stem cells. Nat Methods 11, 847-854.

Yusa, K., Zhou, L. Q., Li, M. A., Bradley, A. and Craig, N. L. (2011). A hyperactive piggyBac transposase for mammalian applications. P Natl Acad Sci USA 108, 1531-1536.

Zhang, D. W., Yang, S. Z., Toledo, E. M., Gyllborg, D., Salto, C., Villaescusa, J. C. and Arenas, E. (2017). Niche-derived laminin-511 promotes midbrain dopaminergic neuron survival and differentiation through YAP. Science Signaling 10.

