## Supplementation for "Spatial cell fate manipulation of human pluripotent stem cells by controlling the microenvironment using photocurable hydrogel"

### Supplementary tables

#### Table S1: List of oligonucleotides

| Amplicon | Forward | Reverse | Ref |
|---|---|---|---|
| Oct4 | CCTGAAGCAGAAGAGGATCACC | AAAGCGGCAGATGGTCGTTTGG | *Origene, HP206340* |
| PAX6 | CTGAGGAATCAGAGAAGACAGGC | ATGGAGCCAGATGTGAAGGAGG | *Origene, HP225517* |
| OTX2 | GGAAGCACTGTTTGCCAAGACC | CTGTTGTTGGCGGCACTTAGCT | *Origene, HP214209* |
| SOX1 | GAGTGGAAGGTCATGTCCGAGG | CCTTCTTGAGCAGCGTCTTGGT | *Origene, HP209017* |
| PBGD | GGAGCCATGTCTGGTAACGG | CCACGCGAATCACTCTCATCT | *Kyle M. Loh et al., 2014* |
| FOXA1 | AAGGCATACGAACAGGCACTG | TACACACCTTGGTAGTACGCC | *Ang et al., 1993; Sasaki and Hogan, 1993* |
| FOXA2 | GGGAGCGGTGAAGATGGA | TCATGTTGCTCACGGAGGAGTA | *Ang et al., 1993; Sasaki and Hogan, 1993* |
| Hhex | CACCCGACGCCCTTTTACAT | GAAGGCTGGATGGATCGGC | *Thomas et al., 1998* |
| SOX17 | CGCACGGAATTTGAACAGTA | GGATCAGGGACCTGTCACAC | *Kanai-Azuma et al., 2002* |
| TBX6 | AAGTACCAACCCCGCATACA | TAGGCTGTCACGGAGATGAA | *Kyle M. Loh et al., 2016* |
| MSGN1 | CGGAATTACCTGCCACCTGT | GGTCTGTGAGTTCCCCGATG | *Kyle M. Loh et al., 2016* |
| CDX2 | GGGCTCTCTGAGAGGCAGGT | CCTTTGCTCTGCGGTTCTG | *Kyle M. Loh et al., 2016* |
| HAND1 | GTGCGTCCTTTAATCCTCTTC | GTGAGAGCAAGCGGAAAAG | *Kyle M. Loh et al., 2016* |
| ISL1 | AGATTATATCAGGTTGTACGGGATCA | ACACAGCGGAAACACTCGAT | *Kyle M. Loh et al., 2016* |
| NKX2.5 | CAAGTGTGCGTCTGCCTTT | CAGCTCTTTCTTTTCGGCTCTA | *Kyle M. Loh et al., 2016* |
| Amplify H2B-tdTomato insert | tcccgggccaatgccagagccagcgaagt | gatcggaattcttacttgtacagctcgtccatgccg | Not applicable |
| Linearize T-2A-EGFP-PGK-Puro | tgtacaagtaagaattccgatcatattcaataaccct | gctctggcattggcccgggattctcttcgac | Not applicable |
| Genomic PCR of left junction | catcttcctgatgattttgtttcttattaatagatacgacaa | gctctggcattggcccgggattctcttcgac | Not applicable |
| Genomic PCR of right junction | tgtacaagtaagaattccgatcatattcaataaccct | ctgtcctcaactatgattttattctgtccttaacag | Not applicable |
| Sequencing left junction | tgagctctgaatatgtgaataatcttttcagtcatct | gctctggcattggcccgggattctcttcgac | Not applicable |
| Sequencing right junction | tgtacaagtaagaattccgatcatattcaataaccct | agttatagtttaacaacacaagagattagctacatatgc | Not applicable |

#### Table S2: List of antibodies

| Target | Manufacturer | Host animal | Dilution |
|---|---|---|---|
| T | R&D, AF2085 | Goat | 500 |
| SOX2 | Abcam, ab79351 | Mouse | 100 |
| TBX6 | R&D, AF4744 | Goat | 500 |
| T | Abcam, ab20680 | Rabbit | 100 |
| SOX17 | R&D, AF1924 | Goat | 40 |
| PAX6 | R&D, AF8150 | Mouse | 1000 |
| SOX1 | Cell signaling, #4194 | Rabbit | 500 |
| CDX2 | Abcam, ab76541 | Rabbit | 500 |
| HAND1 | R&D, AF3168 | Goat | 200 |
| NANOG | Abcam, #4893 | Goat | 500 |
| OCT3/4 | Abcam, ab181557 | Rabbit | 500 |
| Alexa Fluor™ 546 Phalloidin | Invitrogen, A22283 | Not applicable | 500 |
| Alexa Fluor™ 647 Phalloidin | Invitrogen, A22287 | Not applicable | 500 |
| E-cadherin | Takara, ECCD-2 | Mouse | 200 |
| β-Catenin | Sigma, C2206 | Rabbit | 1000 |
| YAP | Santa Cruz Biotechnology, sc-101199 | Mouse | 200 |

#### Table S3: List of fluorescent molecules

| | Manufacturer | Molecular weight | Concentration |
|---|---|---|---|
| Alexa Fluor™ 488 Phalloidin | Invitrogen, A22283 | 1.4kDa | 100ng/ml |
| FITC-dextran 4 | TdB Labs, FD4 | 4kDa | 100ng/ml |
| FITC-dextran 20 | TdB Labs, FD20 | 20kDa | 100ng/ml |
| FITC-dextran 40 | TdB Labs, FD40 | 40kDa | 100ng/ml |

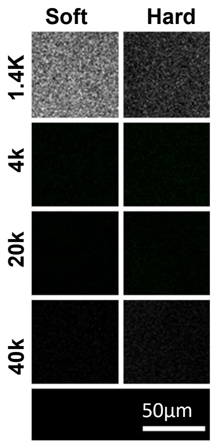


Fig. S1: Fluorescent Diffusion Experiment

Hydrogels were incubated with fluorescent molecules and then observed using laser confocal microscopy to assess the diffusion of fluorescent molecules inside the gel.

Scale bars: 50 µm.

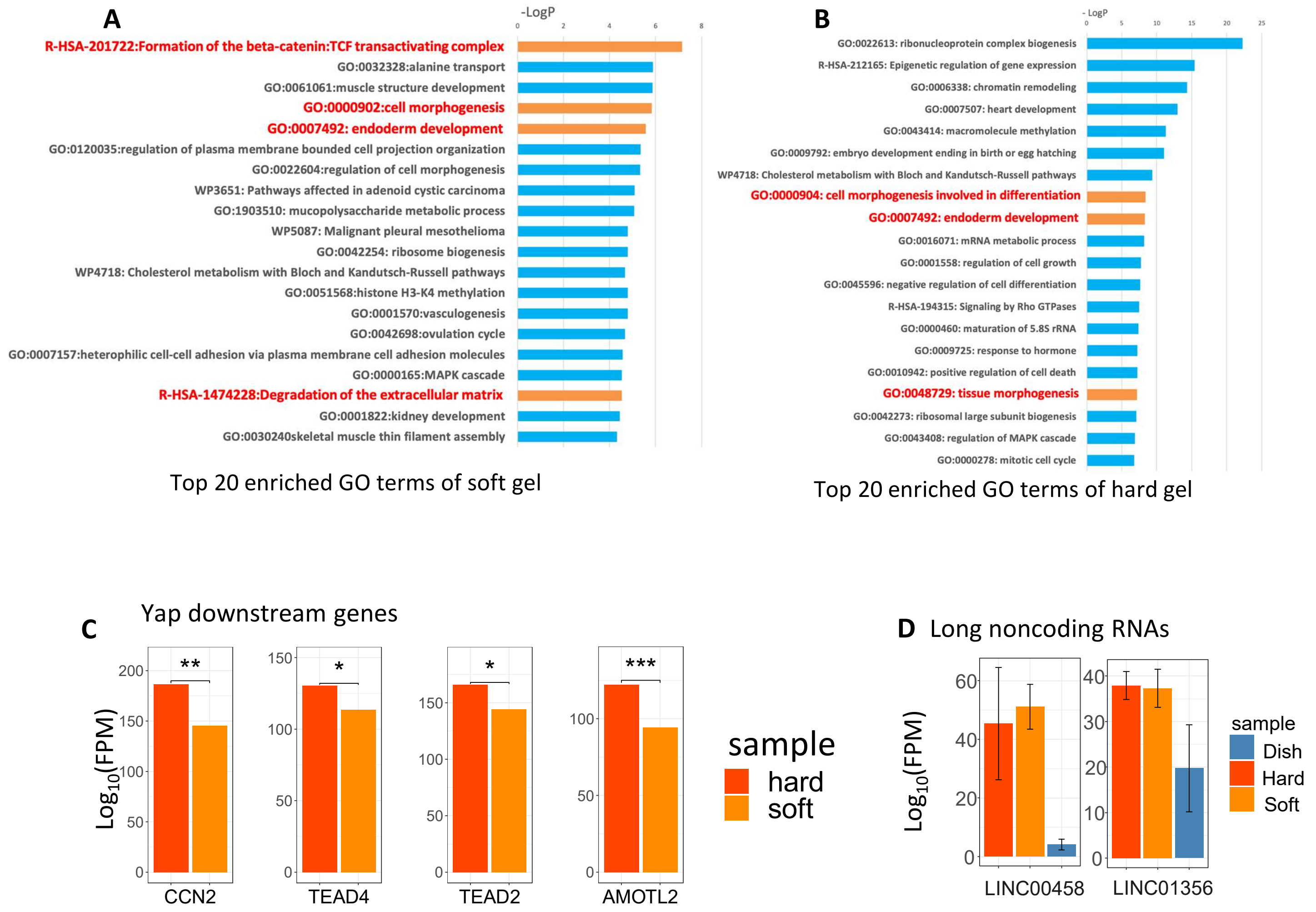


Fig. S2: (A) Top 20 Gene Ontology (GO) Terms (Soft vs. Dish): The top 20 Gene Ontology (GO) terms highlighting the categories of up-regulated genes in the cells on soft gel compared to those on dishes. Terms shown in red are involved in development. (B) Top 20 Gene Ontology (GO) Terms (Hard vs. Dish): The top 20 Gene Ontology (GO) terms highlighting the categories of up-regulated genes in the cells on hard gel compared to those on dishes. Terms shown in red are involved in development. (C) YAP down steam genes Expression Based on RNA-seq: Expression levels of selected genes based on RNA-seq of hPSCs cultured on soft/hard gel. Expression level is shown as TPM (Transcripts Per Kilobase Million). (D) Long non coding RNAs Expression Based on RNA-seq: Expression level is shown as TPM (Transcripts Per Kilobase Million).

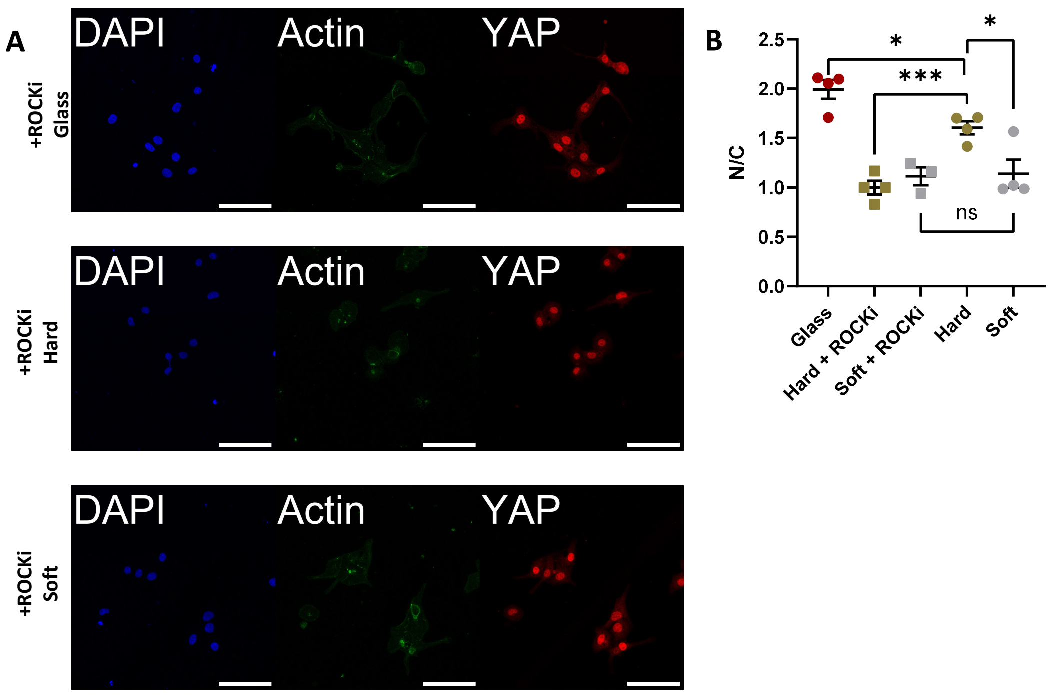


Fig. S3: Immunofluorescent (IF) Staining and YAP Quantification. (A) IF Staining of YAP: Immunofluorescent staining of YAP in cells cultured on a soft/hard gel or glass with 10µM Y-27632. Scale bars: 100 µm. (B) Quantification of YAP Nuclear Localization: YAP nuclear localization was quantified using the Nuclear/Cytoplasmic fluorescent intensity ratio (N/C).

Data obtained from at least 3 biological replicates. Each dot represents a quantification. Mean values, indicated by black lines, and error bars represent standard deviation. Statistical significance was determined using an unpaired, two-tailed t-test: ns - Not significant, *** $P < 0.001$, * $P < 0.1$.

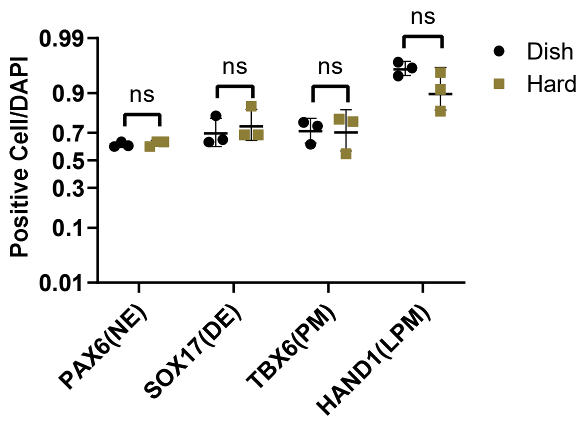


Fig. S4: Differentiation Efficiency Quantification: Positive cell ratio of PAX6 (neural ectoderm), SOX17 (definitive endoderm), TBX6 (Paraxial mesoderm), HAND1 (lateral plate mesoderm).
Data were obtained from at least 3 biological replicates Black lines represent median values. ns: Not significant. Error bars represent standard deviation.

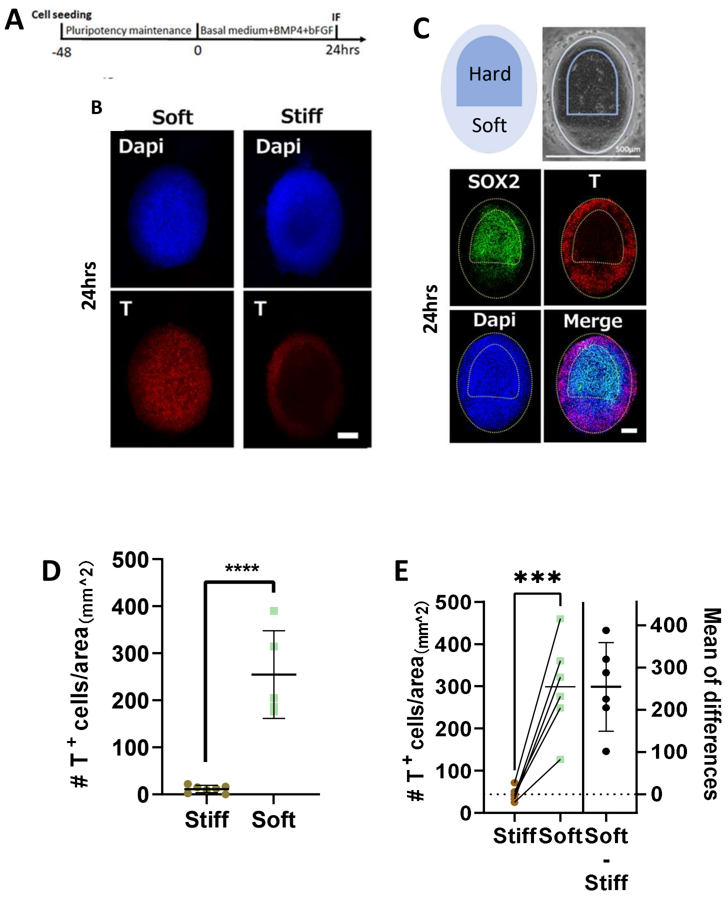


Fig. S5: Manipulating Stem Cell Fate by Adjusting Local Stiffness of the Hydrogel

(A) Differentiation Timetable for Mesoderm Differentiation (B) Immunofluorescence (IF) of 24-hour Mesoderm Differentiation on Soft and Hard Gels Scale bars: 100 µm. (C) IF of 24-hour Mesoderm Differentiation on Gel with Locally Different Stiffness Scale bars: 500 µm in bright filde,100 µm in IF image. (D) Quantification of $T^+$ Number/Area on Soft and Hard Gels (*n*=5) (E) Quantification of $T^+$ Number/Area on Each Gel with Locally Different Stiffness (*n*=6). Dashed line-connected dots represent soft and hard parts from the same patterned gel.

Data were obtained from at least 3 biological replicates. Black lines indicating median values. Error bars represent standard deviation. Statistical significance: **** $P < 0.0001$, *** $P < 0.001$ (unpaired, two-tailed t-test).

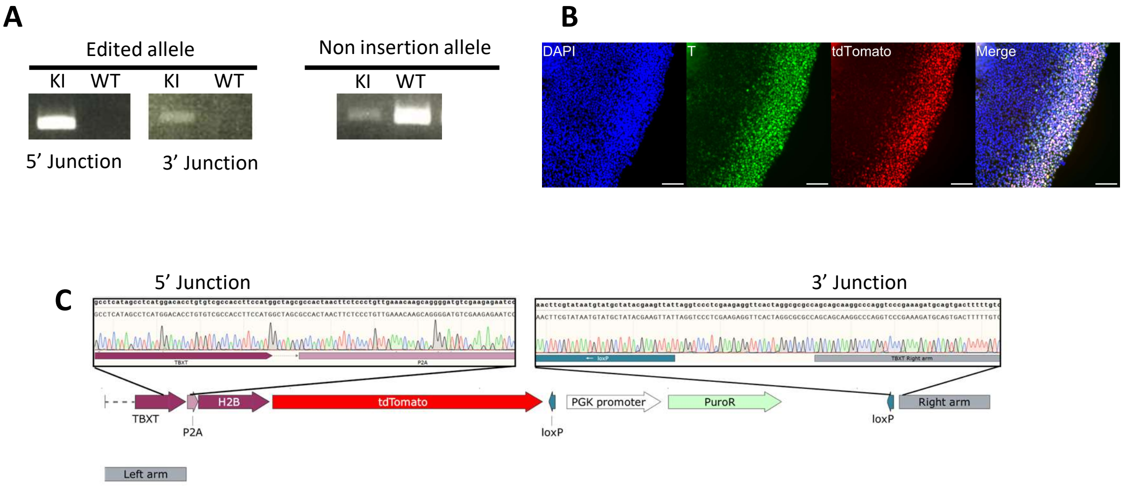


Fig. S6: Verification of Knock-in Reporter T-H2B-tdTomato (A) Genomic PCR of T-H2B-tdTomato: Electrophoresis showing successful heterogeneous integration into the genome. (B) Immunofluorescence (IF) of T-H2B-tdTomato: Differentiated with 10μM CHIR for 24 hours, showing colocalization of tdTomato and T expression. Scale bars: 100 μm. (C) Sanger Sequencing of Junction of Integrated Fragment

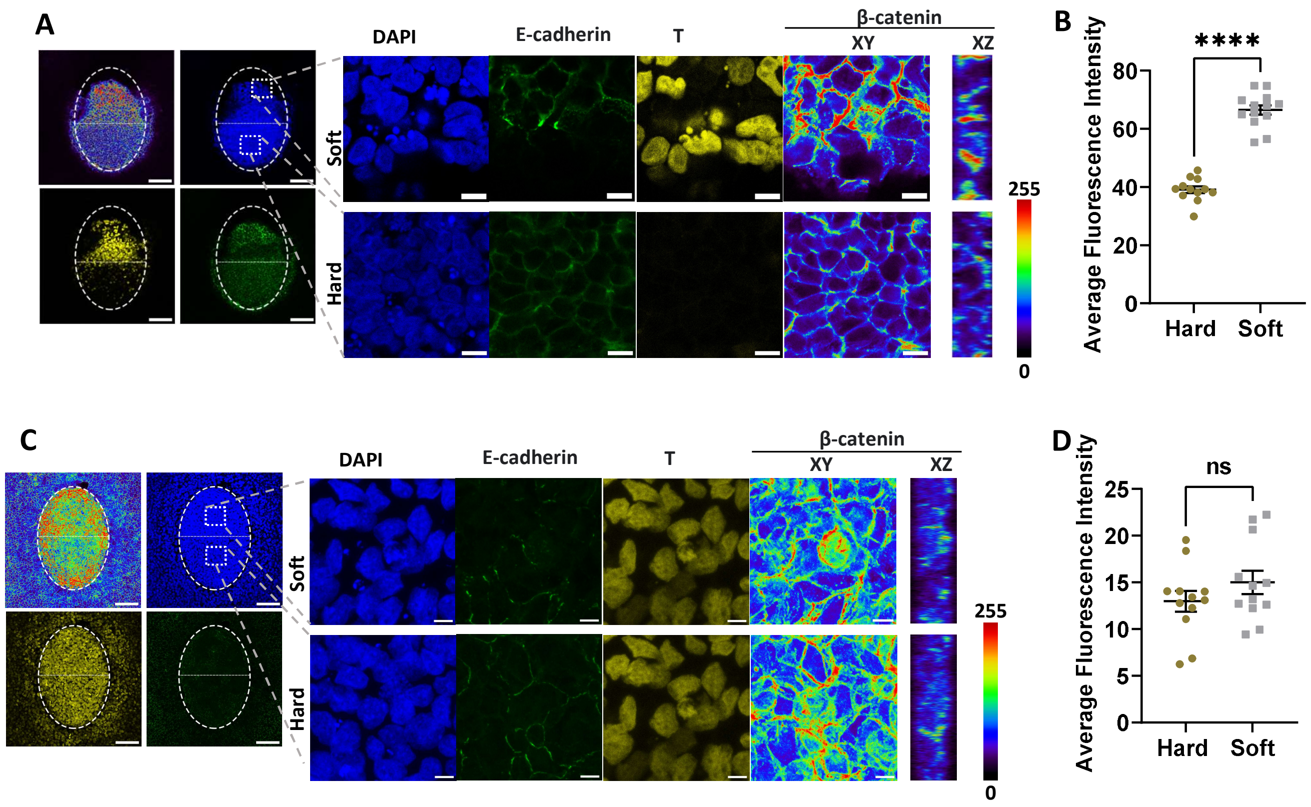


Fig. S7: Stem Cell Fate Control Can Be Achieved with BMP/bFGF but Not CHIR (A) IF Staining (Differentiated with BMP/bFGF): Immunofluorescent staining of hPSCs on a partially soft gel pattern using pulse differentiation, as shown in Fig. 5C. The white dotted oval line indicates the entire gel pattern, with the upper half being soft and the lower half being hard. The white box indicates a magnified view of the indicated region. The β-catenin channel is shown in XY and XZ section views as a heat map, with colors representing fluorescent intensity. Scale bars: 100 µm in the whole gel view and 10 µm in the magnified view. (B) Quantification of β-catenin in BMP/bFGF Differentiation: Quantification of β-catenin fluorescent intensity based on Z sections of IF staining in BMP/bFGF differentiation on a partially soft gel. (*n*=12) (C) IF Staining (Differentiated with CHIR): Immunofluorescent staining of hPSCs on a partially soft gel pattern using pulsed differentiation with CHIR. The white dotted oval line indicates the entire gel pattern, with the upper half being soft and the lower half being hard. The white box indicates a magnified view of the indicated region. The β-catenin channel is shown in XY and XZ section views as a heat map, with colors representing fluorescent intensity. Scale bars: 100 µm in the whole gel view and 10 µm in the magnified view. (D) Quantification of β-catenin in CHIR Differentiation: Quantification of β-catenin fluorescent intensity based on Z sections of IF staining in CHIR differentiation on a partially soft gel (*n*=12).

Data were obtained from at least 3 biological replicates. Each dot represents an individual measurement. The black line represents the mean value. Statistical significance: ns - Not significant **** $P < 0.0001$ (unpaired, two-tailed t-test). Error bars represent standard deviation.

Movie 1: Spatial Cell Fate Control - Live imaging of T-H2B-tdTomato cells undergoing differentiation on a partially soft gel using BMP/bFGF or CHIR. The tdTomato signal is represented in red. The soft area of the gel is highlighted with a yellow circle. The live imaging was recorded at a 1-hour per frame rate.

Movie 2: YAP Inhibition Disrupts Stem Cell Fate Control: Live Imaging of T-H2B-tdTomato Cells Undergoing Differentiation on a Partially Soft Gel using BMP/bFGF in the Presence of 50nM Peptide 17. The tdTomato signal is represented in red, while the soft area of the gel is highlighted with a yellow circle. The live imaging was recorded at a rate of one frame per hour.
